# Early and selective subcortical Tau pathology within the human Papez circuit

**DOI:** 10.1101/2023.06.05.543738

**Authors:** Barbara Sárkány, Csaba Dávid, Tibor Hortobágyi, Péter Gombás, Peter Somogyi, László Acsády, Tim J. Viney

**Affiliations:** Department of Pharmacology, University of Oxford; Oxford, OX1 3QT; UK; Laboratory of Thalamus Research, Institute of Experimental Medicine; Budapest, 1083; Hungary; Department of Anatomy, Histology and Embryology, Semmelweis University; Budapest, 1094; Hungary; Department of Neurology, Faculty of Medicine, University of Debrecen; Debrecen, 4032; Hungary; Department of Pathology, Szt. Borbála Hospital; Tatabánya, 2800; Hungary

## Abstract

The Papez circuit comprises several interconnected brain areas important for spatial navigation and orientation. An early symptom of dementia is disorientation, suggesting that brain regions responsible for providing a sense of direction are adversely affected. We examined *post-mortem* human tissue from cases with no cognitive impairment, mild cognitive impairment, and Alzheimer’s disease. A key part of the Papez circuit, the anterodorsal thalamic nucleus (ADn), contained a high density of misfolded pathological Tau (pTau) at all disease stages, including in control cases. Moreover, pTau preferentially accumulated in calretinin-expressing neurons. At the subcellular level, we detected pTau filaments in ADn cell bodies, dendrites, and in specialized presynaptic terminals. Large vesicular-glutamate-transporter-2-containing terminals from the lateral mammillary nucleus, rather than corticothalamic terminals, preferentially contained pTau, suggesting that Tau crosses specific synapses within the Papez circuit. As the ADn contains a high density of head direction cells, pTau may degrade the processing of orientation signals, explaining why people become disorientated years-to-decades before memory deficits emerge.

## Introduction

Disorientation is emerging as a very early cognitive biomarker predicting future memory deficits associated with Alzheimer’s disease (*1, 2*). Episodic memory impairments are associated with pathology in the entorhinal cortex (EC) and hippocampus (*3, 4*), but pathology in other brain areas may be responsible for the early deficits in spatial orientation. The Papez circuit is a compelling candidate because it comprises several interconnected cortical and subcortical areas that contain head direction cells (*5–7*), including the retrosplenial cortex, anterior thalamus, and mammillary bodies (Fig. 1A). Degradation of head direction signals may explain disorientation.

**Figure 1.**
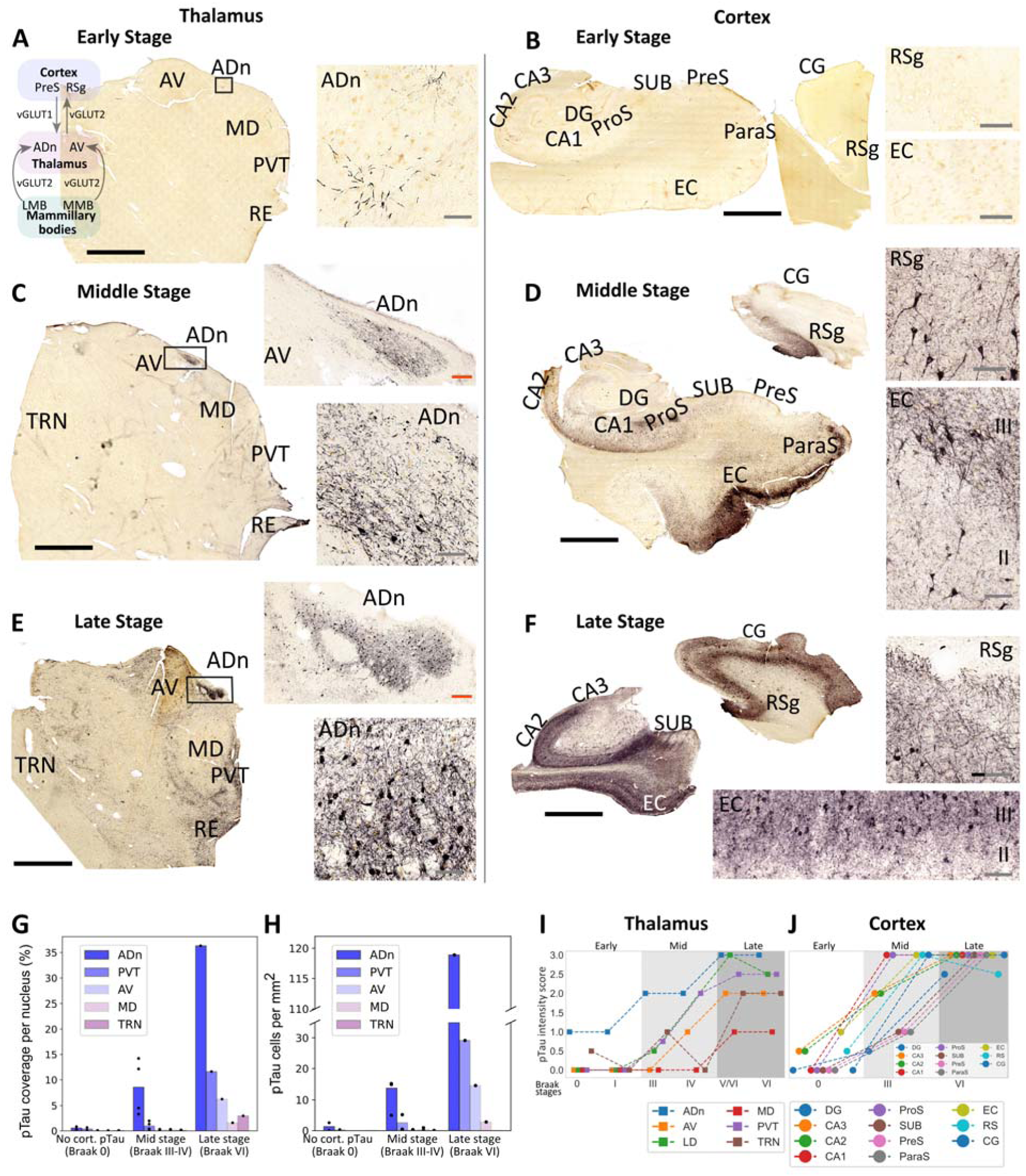
Progression of Tau pathology in the rostral thalamus and the cerebral cortex. (**A-F**) Brightfield images of pTau (AT8 immunoreactivity; HRP-based diaminobenzidine (DAB) end-product) in 50-µm-thick sections of rostral thalamus, hippocampal formation, parahippocampal gyrus, and cingulate gyrus. (A) Braak Stage 0, Case 5. Note pTau+ dendrites in the ADn (right). Inset, schematic of neural circuit. (B) Braak Stage 0, Case 5. Note lack of pTau immunoreactivity in the cortex. (C) Braak stage III, Case 12. (D) Braak stage III, Case 11. (E) Braak stage VI, Case 17. (F) Braak stage VI, Case 17 (cingulate gyrus) and Case 19 (hippocampal formation). (**G-H**) Quantification of pTau coverage (G) and pTau+ cells (H) for early (n=2), middle (n=3), and late (n=1) stage cases. (**I-J**) Scoring of pTau intensity across different thalamic nuclei (n=18 cases) and cortical areas (n=8 cases). Scores: undetectable pTau (0), trace inclusions (0.5), sparse (1), moderate (2), dense (3). Median values are shown (see Table S4 for individual datapoints). Abbreviations: ADn, anterodorsal nucleus; AV, anteroventral nucleus; MD, mediodorsal thalamic nucleus; PVT, paraventricular thalamic nucleus; RE, reuniens nuclear complex; TRN, reticular thalamic nucleus; DG, dentate gyrus; ProS, prosubiculum; PrS, presubiculum; ParaS, parasubiculum; EC, entorhinal cortex; CG, cingulate gyrus; RSg, granular retrosplenial cortex; LMB, lateral mammillary nucleus; MMB, medial mammillary nucleus. Scale bars: black, 4 mm; red, 250 µm; gray, 80 µm. See also Figures S1-S4 and Tables S1-S4.

The anterior thalamic nuclear group, located within the rostral part of the thalamus, forms a key subcortical node of the Papez circuit. It comprises anterodorsal (ADn), anteroventral (AV), and anteromedial nuclei, which receive glutamatergic input from different regions of the mammillary bodies, and project to key cortical areas important for spatial navigation, orientation, and episodic memory (Fig. 1A). Lesions of the anterior nuclear group impair spatial memory in rodents (*8–10*), and degeneration in humans causes memory deficits (*11*). Earlier reports have shown pathological misfolded forms of Tau (pTau), extracellular amyloid deposits, and cell loss in the anterior nuclear group (*12, 13*). Importantly, the ADn has been shown to be the most strongly affected by pTau inclusions, and it contains the highest density of head direction cells (*12, 14, 15*), indicating a potential role for the ADn in driving Alzheimer’s disease-related impairments, especially disorientation.

The apparent ‘prion-like’ spread of pTau is associated with a wide range of neurodegenerative diseases including Alzheimer’s disease (*12, 14, 16*). However, the subcellular compartments, such as axons and dendrites, containing pTau and the temporal progression of pathology have not been fully determined within intact human brain samples, with descriptions typically limited to neurofibrillary tangles and neuropil threads. Compared to the vast and layered cerebral cortex, adjacent thalamic nuclei (or even subfields of the same nucleus) exhibit markedly different cortical and subcortical connectivity (*17–20*). The organization of the Papez circuit thus provides a unique opportunity for understanding the cellular and subcellular spread of pTau.

## Results

### Strong and early vulnerability of the anterodorsal thalamic nucleus to Tau pathology

We examined the rostral thalamus at early (n=9), middle (n=5), and late (n=4) ‘Braak Tau stages’ of disease progression (see Methods) (*14*) and compared these with the hippocampal formation, parahippocampal gyrus, and cingulate gyrus (Fig. 1A-F, S1 and S2; Table S1, S2, S3 and S4). We quantified pTau coverage and pTau-immunoreactive (pTau+) cells using the AT8 antibody, which recognizes phosphorylated serine 202 and threonine 205 residues of Tau (*21*), in 50-µm-thick sections from perfusion-fixed human brains using a pixel classifier (Fig. 1G and H, S3; Table S1; Methods). We confirmed these data with a pTau-intensity scoring system in a larger number of samples comprising the perfusion-fixed sections and 10-µm-thick formalin-fixed paraffin-embedded (FFPE) sections (Fig. 1I and J; Table S4; n=18 cases).

We observed pTau+ cells and processes restricted to the ADn in ‘controls’ (stage 0; n=3/4 cases; coverage=0.56%, pTau+ cells=1.29 mm^-2^, intensity score=1; Fig. 1A, G, H and I, 2A). In contrast to the ADn, at early stages (defined here as Braak stages 0-I), other rostral thalamic nuclei, even those adjacent to the ADn such as the AV nucleus, distinctly lacked pTau (Fig. 1A and 2A; Table S3 and S4). Among the postsynaptic targets of the ADn (*22, 23*), both the EC and granular retrosplenial cortex (RSg), but not the presubiculum (PreS), exhibited sparse pTau (EC score=1, n=3 cases; RSg score=0.5, n=2 cases; Fig. 1B and J; Table S4).

**Figure 2.**
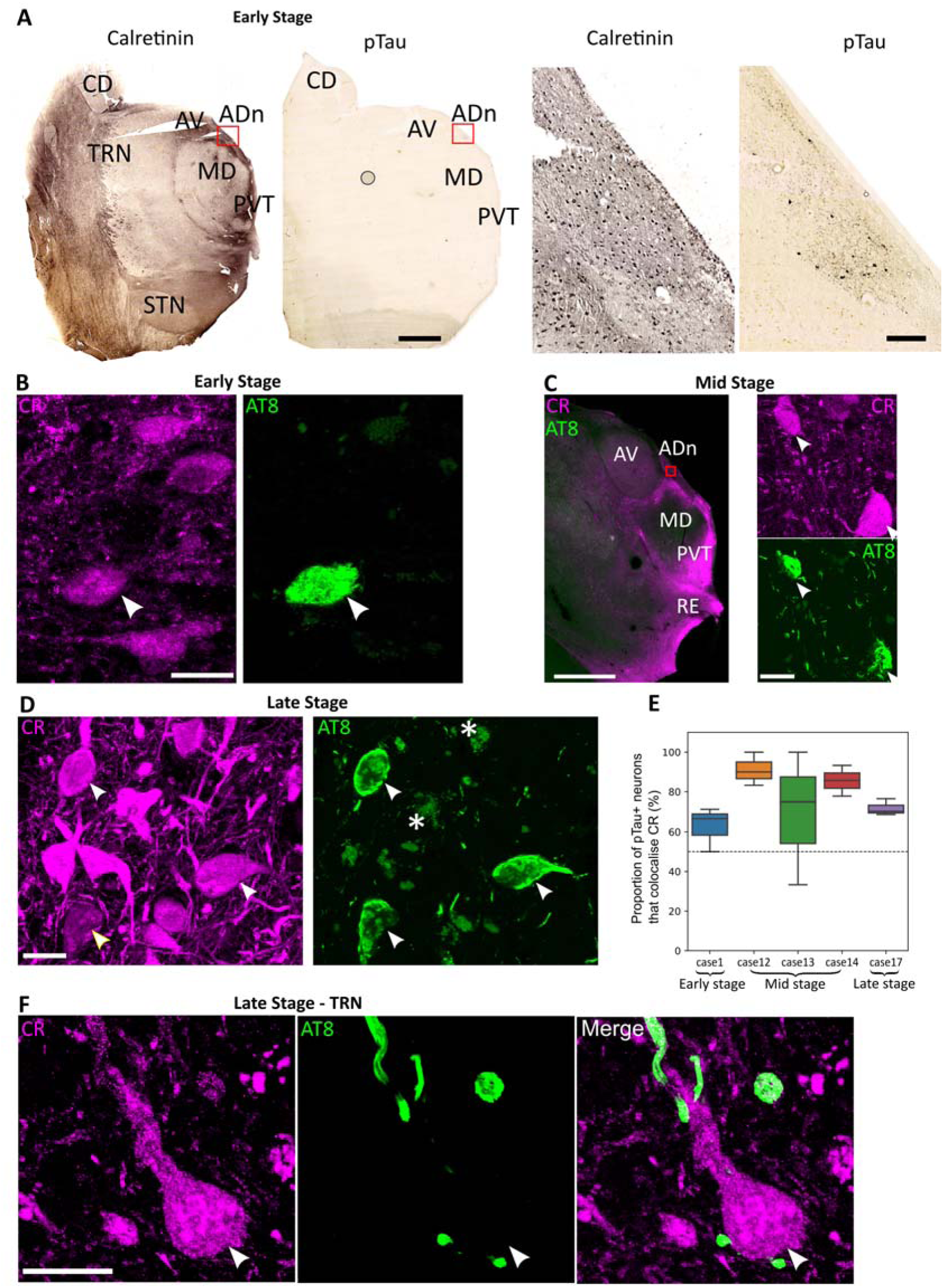
pTau-containing neurons in the anterodorsal thalamic nucleus express calretinin. (**A**) Brightfield images of the rostral thalamus at Braak Stage 0 (Case 1, FFPE, 10-µm-thick section). Left, adjacent sections immunoreacted for calretinin (CR) and AT8 (pTau) visualized with HRP-based DAB reaction end-product. Right, enlarged view of the ADn (red boxed region) revealing CR+ and pTau+ neurons. (**B-D**) Subpopulations of calretinin (CR) immunoreactive neurons (magenta) colocalize pTau (AT8, green; white arrowheads) in the ADn. (B) Early stage (Case 1, FFPE sample), confocal z-projection (5.7 µm thick). (C) Middle stage (Case 12, perfusion-fixed, 50-µm-thick section). Left, widefield fluorescence. Right, detail CR and AT8 colocalization in the ADn from an adjacent section (red boxed region), confocal z-projection (3.7 µm thick). (D) Late stage (Case 17, perfusion-fixed, 50-µm-thick section), confocal z-projection (3.9 µm thick). Faded green signal is lipofuscin (e.g. asterisks). (**E**) Box plots of the percentage of AT8 immunoreactive cells in the ADn that were immunopositive for CR. (**F**) In the TRN, a CR+ neuron in close apposition to pTau+ boutons lack pTau (arrowhead, soma). Confocal z-projection, 2.2 µm thick (C). Scale bars: 4 mm (A left, C left); 250 µm (A right); 100 µm (C right); 20 µm (B, D, F). Abbreviations: CD, caudate nucleus; STN, subthalamic nucleus. See also Tables S1-S3.

At middle stages (Braak stages III-IV), the ADn contained moderate levels of pTau (coverage=8.54%, pTau+ cells=11.65 mm^-2^, intensity score=2; Fig. 1C, G, H and I, S1; Table S3 and S4), whereas other rostral thalamic nuclei showed sparse pTau (AV: coverage=0.15%, pTau+ cells=14.49 mm^-2^, score=0.5; paraventricular nucleus (PVT): coverage=0.96%, pTau+ cells=1.91 mm^-2^, score=1.35; Fig. 1C, G, H and I, S1; Table S3 and S4). In the cortex, pTau was observed at moderate-to-intense levels, with the RSg, EC and prosubiculum (ProS) showing intense pTau immunoreactivity; the PreS displayed sparse immunolabeling at this stage (RSg score=3; EC score=3; ProS score=3; PreS score=1; Fig. 1D and J; Table S4).

At late stages (Braak stages V-VI), which had Alzheimer’s disease (Table S1 and S2) (*14*), intense pTau was observed in the ADn, with a high density of pTau+ cells (coverage= 36.31%, pTau+ cells=118.89 mm^-2^, intensity score=3; Fig. 1E, G, H and I, S2; Table S3 and S4). The laterodorsal nucleus (LD) showed a high pTau density in the late stages, matching that of the ADn (Fig. 1I and S2; Table S4). The PVT had a similar pTau coverage to middle-stage ADn (11.62%, score=; Fig. 1C, E, G and I, S2; Table S3 and S4) and had the highest pTau+ cell count after the ADn (29.13 cells mm^-2^). The reuniens nuclear complex (RE) was similarly affected (score=2). In contrast, the AV had lower coverage and pTau+ cell counts (coverage=6.28%, pTau+ cells=14.49 mm^-2^, intensity score=2) followed by the mediodorsal nucleus (MD), which remained relatively sparse compared to the other examined nuclei (coverage=1.62%, pTau+ cells=2.84 mm^-2^, score=1; Fig. 1E, G, H and I; Table S3 and S4). The GABAergic thalamic reticular nucleus (TRN) lacked pTau immunopositive cell bodies; only axons and axon terminals were pTau+ (coverage= 2.98%, pTau+ cells=0 mm^-2^, score=2 Fig. 1E, G, H and I, S2, Table S3 and S4), consistent with published data (*24*). In the cortex, pTau severely affected each examined area (Fig. 1F and J; Table S4), consistent with previous studies (*14*).

In addition to neuronal pTau, we observed pTau-immunopositive (pTau+) ‘coiled bodies’ in the ADn (Fig. S4A). Based on their size (∼10 µm) and shape, we suggest that coiled bodies are localized to oligodendrocytes, which are typically overlooked in Alzheimer’s disease (*25, 26*). We also detected pTau+ astrocytes, which are associated with both ageing and Alzheimer disease (*27, 28*). Interestingly, we could find pTau+ astrocytes of varying shape and size, even within the same brain regions of the same cases (Fig. S4B, Table S2). This suggests that pTau can accumulate in a variety of different kinds of astrocytes (*27*). These results reveal that distinct nuclei of the rostral thalamus are affected early on by pTau, with the ADn consistently having the highest pTau density and pTau+ cells across all stages (Fig. 1A, C, E, G, H and I, 2A).

### Calretinin-expressing neurons accumulate pTau in the rostral thalamus

We noticed that thalamic nuclei vulnerable to pTau were in calretinin (CR)-enriched regions (Fig. 2A, S1, S2) (*29*) and hypothesized that CR+ neurons were sensitive to accumulating pTau. We performed double immunolabeling with CR and AT8 (for pTau) and observed colocalization in neurons within the ADn, PVT, and RE (Fig. 2A-D). In the TRN, CR-enriched neurons lacked pTau (Fig. 2F), consistent with the distinct lack of pTau+ TRN cell bodies (Fig. 1H). Since the ADn contained pTau+ neurons even in control cases (stage 0; Fig. 1A and H, 2A), we tested whether CR+ neurons were affected at early, middle and/or late stages. Even at Braak stage 0, CR was detected in the majority of pTau+ neurons (64.3% CR+, n=1 case; Fig. 2A, B, and E). In the middle stage, a large proportion of pTau+ cells were CR+ (81.1%; n=3 cases; Fig. 2C and E), and in the late stage, 71.1% were CR+ (n=1 case; Fig. 2D and E). In conclusion, CR-expressing neurons were affected very early on, and at every stage, the majority of pTau immunopositive cells were CR+ in the ADn.

### Subcellular distribution of pTau in the anterodorsal thalamus

After establishing that ADn neurons were especially vulnerable to pTau, we investigated the subcellular distribution of pTau in order to reveal how it spreads. To define synaptic structures at different stages of Tau pathology, we examined ultrathin (∼50-70 nm) sections of the ADn. We were able to obtain electron microscopic samples from 5 cases (Cases 3, 4, 12, 17; Table S1 and S2); 3 cases were appropriately preserved for quantitative analysis (Fig. 3A-D; Braak stages 0, III, and VI). We identified two main types of synaptic boutons with asymmetric synapses: large ∼1-8 µm boutons (Fig. 3C, Fig. S5A-C), consistent with presynaptic axon terminals from the mammillary body (*30*), and small <∼1 µm diameter boutons (Fig. 3D, Fig. S5A-C), resembling ‘classical’ cortical presynaptic terminals(*31*). Some presynaptic boutons from stage 0 (n=20/70) and stage VI (n=31/103) had a highly electron opaque (‘dark’) appearance, ranging from a homogeneous state to others with recognisable vesicles and mitochondria, but all showing collapsed, scalloped forms (Fig. S6A). This may indicate degeneration of certain nerve terminals, and/or be a sign of selective vulnerability to *post-mortem*/fixation conditions (*32, 33*); these terminals were omitted from our quantification.

**Figure 3.**
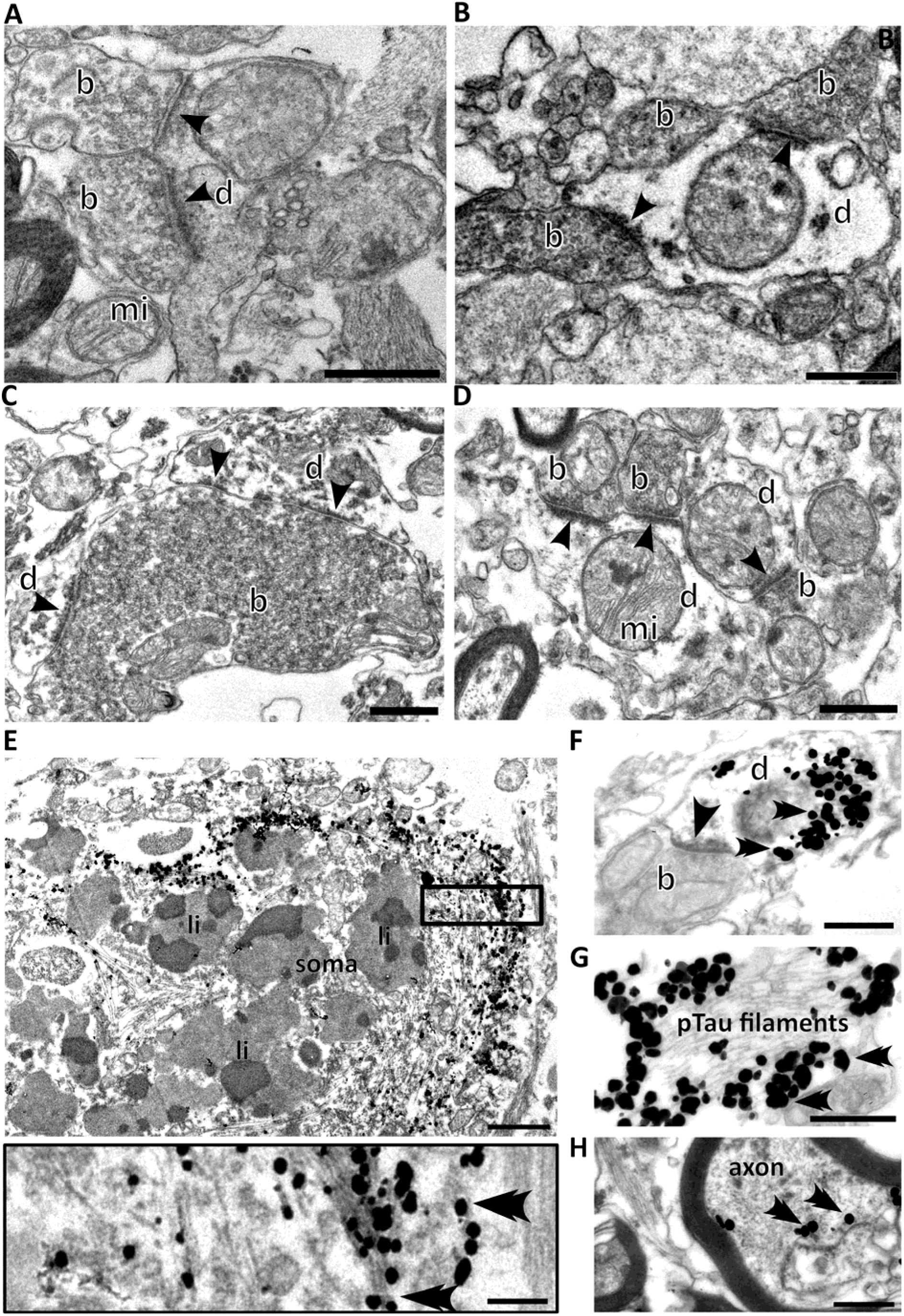
Electron micrographs of synaptic elements and Tau filament-bearing structures in the human anterodorsal thalamic nucleus. (**A-D**) Unlabeled synaptic boutons (b) and dendrites (d) forming synaptic junctions (arrowheads): (A, B) early stage (Braak stage 0, Cases 3 and 4); (C) middle stage (Braak stage III, Case 12); a large bouton containing a high density of synaptic vesicles forms multiple synapses with its postsynaptic partners; (D) late stage (Braak stage VI, Case 17), small terminals form junctions (arrowheads) with small caliber dendrites. (**E-H**) Immunolabeling for pTau (AT8, gold-silver particles, all from Case 17, Stage VI): (E) localization of pTau in a cell body containing lipofuscin (li); inset, detail of the boxed region showing immunolabeled pTau filaments (double arrowheads mark gold-silver particles); (F) a thin, p-Tau+ dendrite receiving an asymmetric synapse (arrowhead) from a small bouton (b); (G) high power image of paired helical filaments associated with silver intensified immunogold particles demonstrating pTau immunoreactivity; (H) pTau immunolabeling in a myelinated axon. Scale bars: 0.5 µm (A-H), 0.25 µm (E inset). See also Figures S5 and S6, and Tables S1 and S2.

To identify subcellular pTau, we first examined cell bodies in the ADn, which contained abundant filaments (Fig. 3E). These resembled filaments previously found in the cortex of tauopathies including Alzheimer’s disease (*16, 34, 35*). We visualized pTau with silver-enhanced immunogold particles, and observed that pTau was specifically associated with the intracellular filaments (Fig. 3E), thus unequivocally demonstrating the association of pTau with the originally described paired helical filaments (*36*) at the ultrastructural level. Cell bodies also contained abundant lipofuscin (Fig. 3E and 2D). Filaments immunolabeled for pTau were also localized to dendrites (Fig. 3F, S6B-D), and could be observed in large bundles (>1 µm) (Fig. 3G, S6D). Filament bundles were immunolabeled predominantly on the cytoplasmic surface, most likely due to reagents not penetrating into the bundle (Fig. 3G). We also detected pTau in myelinated axons (Fig. 3H). Given that pTau was localized to a variety of subcellular domains, we next investigated whether pTau can also be associated with axon terminals in the ADn.

### Subcortical vesicular transporter 2-expressing presynaptic terminals preferentially contain pTau

Large presynaptic terminals of subcortical origin contain vesicular glutamate transporter 2 (vGLUT2) (*20*). We observed strongly overlapping distributions of vGLUT2 and AT8 immunoreactivities at the light microscopic level, especially in the ADn, RE, PVT, and internal medullary lamina (Fig. 4A). The overlapping vGLUT2 and AT8 distributions suggested that vGLUT2 may be related to pTau. When we examined ultrathin sections immunoreacted for both vGLUT2 and AT8, we discovered that pTau was localized to vGLUT2+ boutons (Fig. 4B, S6B). Whereas some vGLUT2+ boutons showed no signs of abnormalities (Fig. 4C), others were degenerating (Fig. 4D), similar to the unlabeled material (Fig. S6A). The degenerating vGLUT2+ boutons had clumped mitochondria (Fig. 4D). Many of these boutons contained large (80-100 nm) double-walled vesicles (Fig. 4D) (*34*), consistent with autophagy or the packaging and/or potential release of different forms of Tau (*37, 38*). Not all vGLUT2-positive degenerating boutons displayed detectable pTau, at least in the sections that we examined. In some degenerating boutons which were immunoreactive for both vGLUT2 and pTau, pTau-positive bundles of filaments occupied a large proportion of the volume crowding out vesicles (Fig. 4B and E), which may cause impairments in neurotransmission. We also observed synaptic partners consisting of presynaptic vGLUT2+ boutons and postsynaptic dendrites that *both* contained pTau (Fig. 4F, S6B), suggestive of transsynaptic spread between the mammillary bodies and ADn (Fig. 1A). The size distribution of pTau-positive terminals also confirmed that large presynaptic terminals are preferentially affected by pTau-pathology (Fig. S5A-C).

**Figure 4.**
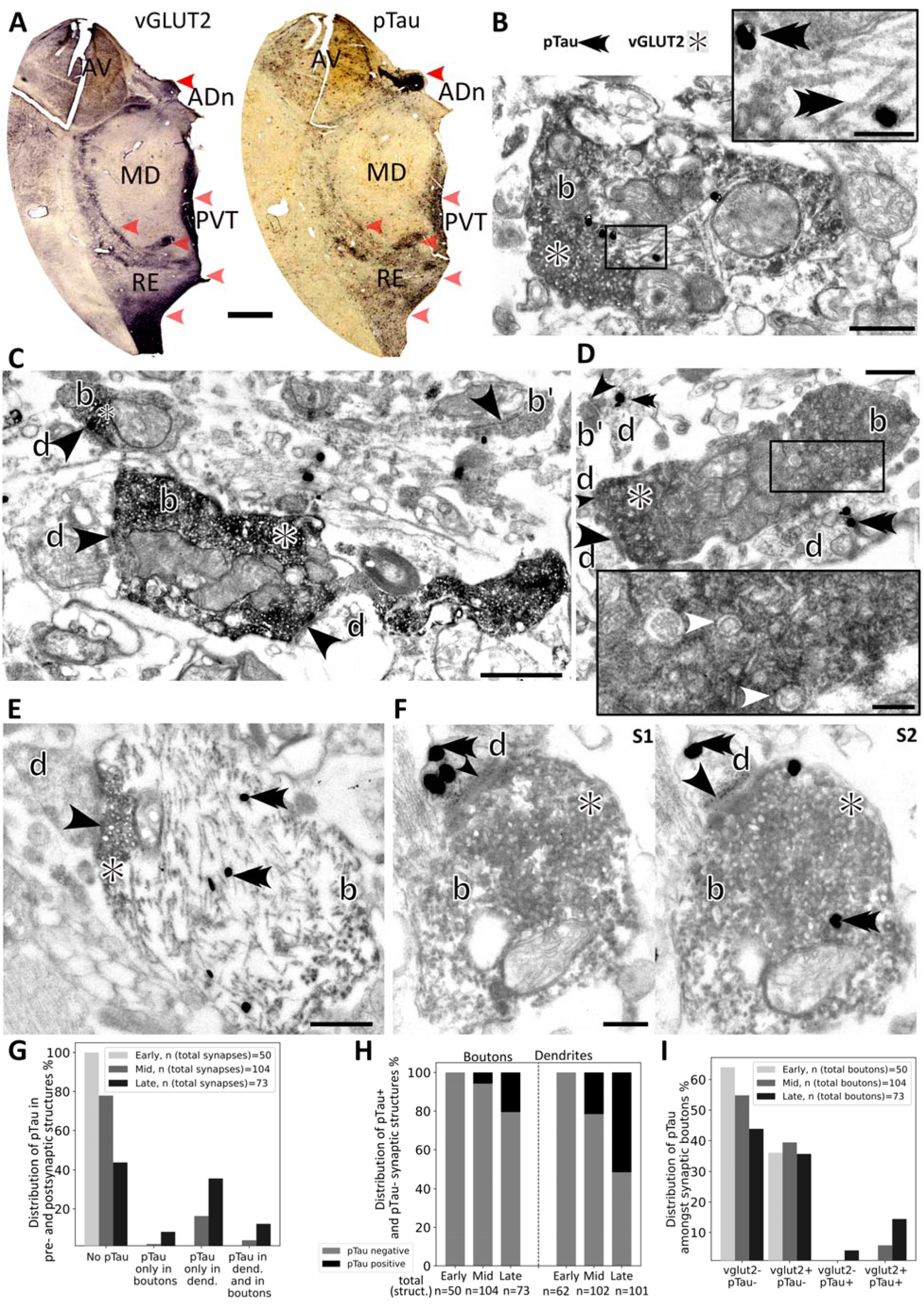
Localization of pTau to vGLUT2-expressing presynaptic terminals. (**A**) Brightfield microscope images of rostral thalamus (serial sections, late stage, Case 17) reacted for vGLUT2 (left) and pTau (AT8, right); HRP-based diaminobenzidine end-products (DAB). The distributions correlate (red arrowheads). (**B-F**) Electron micrographs showing pTau filaments (AT8 immunolabeling, e.g. double arrowheads mark gold-silver particles) and vGLUT2-immunolabeled boutons (DAB) from the ADn of Case 17. Abbreviations: b, bouton; d, dendrite. Black arrowheads mark synaptic junctions. (B) A vGLUT2-immunopositive (vGLUT2+) bouton containing pTau filaments in the middle. Inset, higher magnification of helical pTau filaments from boxed region. (C) Two vGLUT2+ boutons (DAB) and a vGLUT2-bouton (b’, top right) lacking pTau, a silver particle is adjacent to the bouton. (D) A large vGLUT2+ terminal (b) containing double-walled vesicles and clumped mitochondria. Inset, detail of boxed region showing vesicles (e.g. marked by white arrowheads). A small bouton (b’, top left) forms a synapse with a pTau immunopositive dendrite. (E) A large vGLUT2+ bouton (note DAB end product on vesicles at the synapse) containing dense pTau filaments. (F) Electron micrographs of sequential sections (S1, S2) showing pTau in both the vGLUT2+ bouton and the postsynaptic dendrite. Double arrowheads mark examples of gold-silver particles; asterisks label dark DAB product around vGLUT2+ vesicles. (**G**) Proportion of synapses (%) in the ADn with or without pTau in the presynaptic bouton and/or postsynaptic dendrite in early, middle, and late Braak stages of tauopathy. (**H**) Proportions of synaptic boutons (%, left) and dendrites (right) containing pTau at the different stages. (**I**) Proportion of synaptic boutons (%) with or without vGLUT2 and pTau immunoreactivities at the different stages. Scale bars: 2mm (A), 0.5 µm (B,D,E), 1 µm (C), 0.25 µm (F), 150 nm (B and D insets). See also Figures S5 and S6, and Tables S1 and S2.

### The distribution of presynaptic and postsynaptic pTau suggests transsynaptic spread

Given the observation of pTau in both presynaptic terminals and postsynaptic dendrites (Fig. 3F, 4B, D-F, S6B-D), we quantitatively characterized how the synaptic distribution of pTau changed across different Braak stages, examining 543 presynaptic boutons and postsynaptic dendrites, each of which was followed over several serial sections. In the early stage (Braak stage 0), despite pTau being detectable at the light microscopic level (Fig. 1A and 2A), all sampled boutons and dendrites lacked pTau (n=50/50 boutons, n=62/62 dendrites; Case 4; Fig. 4G and H). In the middle stage (Braak stage III), 5.8% of boutons (n=6/104) and 21.6% of dendrites (n=22/102) were pTau+ (Case 12; Fig. 4G and H). In this stage the proportion of synapses in which *both* the presynaptic boutons and the associated postsynaptic dendrites contained pTau was 3.9% (n=4/104; Fig. 4F and G, S6B). In the late stage (Braak stage VI), the proportion of affected boutons and dendrites greatly increased: 20.6% of boutons (n=15/73) and 51.5% of dendrites (n=52/101) contained pTau (Case 17; Fig. 4B, D-H). Furthermore, 12.3% (n=9/73) of synapses consisted of *both* pTau+ boutons and pTau+ dendrites (Fig. 4F and G). These data demonstrate that the proportions of both the presynaptic and postsynaptic elements containing pTau increase with Braak stage.

Finally, we examined the relationship between presynaptic vGLUT2 and pTau across stages. In the early stage, we found vGLUT2+ boutons (Fig. S5E) but did not detect pTau (Fig. S5A). But by the middle stage, from a total of 104 synaptic boutons, 5.8% (n=6) were both vGLUT2 and pTau double immunopositive (Fig. 4I), whereas none of the vGLUT2 immunonegative boutons (n=57) were pTau+. In other words, 100% of pTau+ boutons were vGLUT2+ (n=6) and 12.8% of vGLUT2+ boutons (n=47) were pTau+, supporting the hypothesis of selective vulnerability of subcortical vGLUT2+ synaptic terminals. Filamentous contacts with postsynaptic structures, known as puncta adherentia, are associated with mammillothalamic terminals(*20*). We identified puncta adherentia between vGLUT2+ boutons and postsynaptic dendrites containing pTau (Fig. S6B). Moreover, the small corticothalamic boutons lacked pTau (Fig. S5B and E), which indicates that pTau in the ADn is unlikely to have spread anterogradely from the cortex.

In the late stage, out of a total of 73 synaptic terminals, an even higher proportion showed vGLUT2 and pTau colocalization (16.4%; n=12; Fig. 4I), i.e., *∼80%* of pTau+ boutons (n=15) were immunopositive for vGLUT2. And of all vGLUT2+ boutons (n=38), 31.5% were pTau+. The data on the colocalization of vGLUT2 and pTau is even likely to be an *underestimate*, given that large ‘dark’ boutons are likely to be degenerating mammillothalamic terminals (Fig. S5F, S6A), and we only sampled relatively few sections from each terminal. The above results suggest that vGLUT2+ boutons are strong candidates for the transsynaptic spread of pTau between postsynaptic ADn neurons and presynaptic mammillary body neurons within the Papez circuit (Fig. 5).

**Figure 5:**
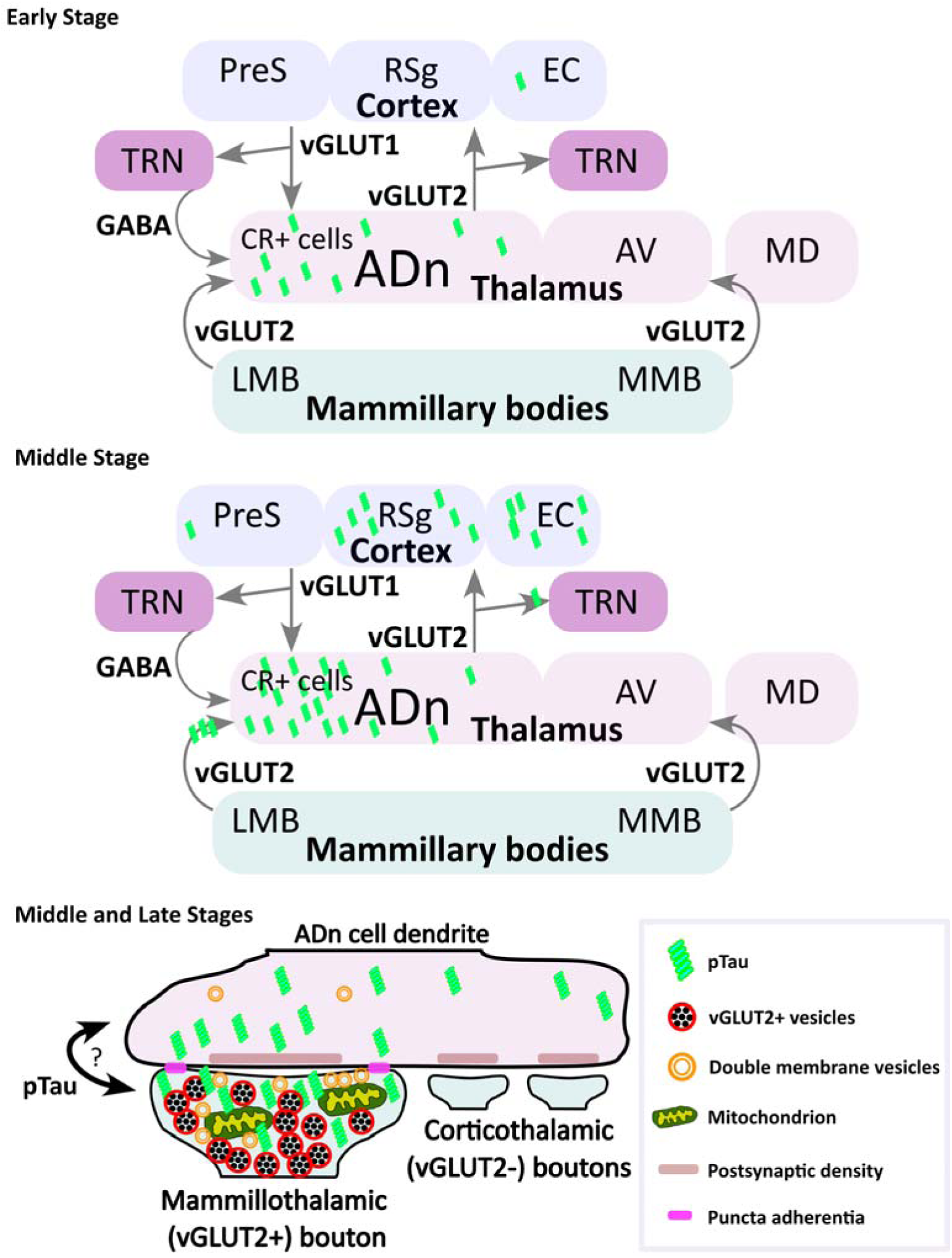
Schematic of pTau transsynaptic spread involving vGLUT2+ terminals in the ADn. Scheme summarizing the data. According to our hypothesis, in the early stage (top), pTau (green) initially accumulates in CR+ cells of the ADn. By the middle stage, pTau has accumulated in 2 of 3 cortical areas postsynaptic to the ADn: the EC and RSg, but not the PreS. In the thalamus, pTau is present in axons innervating the TRN and in mammillothalamic (vGLUT2+) terminals innervating the ADn. The GABAergic cells in the TRN are not invaded by pTau, nor are the corticothalamic terminals in the ADn. Bottom, detail of pTau within the ADn at the middle stage (also applies to the late stage). Transsynaptic spread between the large vGLUT2+ mammillothalamic boutons and ADn dendrites may be retrograde or anterograde, potentially mediated through puncta adherentia (see Fig. S6B). Puncta adherentia are not found between small corticothalamic boutons and postsynaptic dendrites, and pTau was not observed in these presynaptic terminals at either middle or late stages, despite pTau in presynaptic cortical areas. See Discussion for further details.

## Discussion

### Significance of ADn Tau pathology

Untangling which brain regions are first affected in neurodegenerative diseases may facilitate earlier diagnosis and enhance treatment options. We found that a key component of the Papez circuit that contains a high density of head direction cells, the ADn, accumulated pTau very early on, and that affected neurons were detected even in control cases lacking significant neocortical and hippocampal pathology (Braak stage 0). Calretinin-expressing neurons were vulnerable at all disease stages. Given the consistently high pTau density and pTau+ cell counts that we observed in the ADn at all stages, we suggest that the human ADn is a ‘hub’ for early-forming Tau pathology. Using high-quality perfusion-fixed tissue and immunogold labeling, we localized, for the first time, pTau to intact presynaptic boutons and postsynaptic dendrites, with paired helical filaments resembling those described previously in cases of Alzheimer’s disease and other tauopathies (*16, 27, 34, 36, 39*). Surprisingly, despite the classical hypothesis of pTau spreading via corticocortical pathways originating in the EC, pTau was not found in corticothalamic boutons in the ADn. Instead, pTau was preferentially localized to large vGLUT2-expressing subcortical terminals. This suggests an alternative or parallel major subcortical glutamatergic pathway necessary for spatial orientation is driving the spread of pTau in relation to the emergence of Alzheimer’s disease.

The thalamic ADn, AV, and AM are commonly referred to as the anterior nuclear group; lesions impair spatial reference and working memory in rats (*8, 9, 40, 41*), and thalamic infarctions (especially those that include the mammillothalamic tract) are associated with amnesia in humans (*42*). Also, in mice, selective disruption of the ADn impairs spatial working memory (*43*). The anterior nuclear group shares cortical targets through direct projections to the RSg, PreS, and EC (*22, 23*). The AV and AM receive inputs from multiple cortical areas including the EC, yet the ADn only receives input from the dorsal part of the RS and not the EC (*44, 45*). If pTau had spread directly from the EC to the anterior nuclear group, we would have expected to find early pTau in the AV, but it was sparse even at the middle stage. Another key difference between the anterior thalamic nuclei is that the ADn receives glutamatergic input from the lateral mammillary nucleus, whereas the AV and AM receive input from the medial mammillary nucleus (*46, 47*) (Fig. 1A, Fig. 5).

The large vGLUT2-containing axon terminals arising from the lateral mammillary nucleus are thought to act as ‘drivers’, releasing glutamate in response to dynamic changes in the pattern of sensorimotor inputs evoked by stimuli, such as changes in head direction or shifts in gravity (*15, 20, 48, 49*). Postsynaptically, ADn neurons will be strongly depolarized leading to high-frequency firing within the receptive field (*15*). The lateral mammillary nucleus and ADn are more severely affected by Tau pathology compared to the medial mammillary nucleus, AV, and AM (*12, 50, 51*). It is likely that pTau accumulating within axon terminals and neurons in the ADn will disrupt receptive fields (eg. head direction tuning, angular velocity) and firing rates, thereby decreasing the information content provided to postsynaptic neurons in the RSg, PreS, and EC (*22, 23*). This might cause early and progressive deficits in the awareness of orientation and a resulting increase in the probability of losing balance. Our findings may explain the early impairments in spatial navigation and orientation, memory deficits, and an increased number of falls in people that go on to develop Alzheimer’s disease (*52–54*).

### Propagation of pTau involving vGLUT2-containing terminals

Nuclei adjacent to the ADn such as the AV and MD were relatively resistant to pTau, even in late stages, suggesting that in the thalamus, propagation of pTau is facilitated via specific synapses and circuits rather than geometrical proximity. Our data are consistent with the following hypothesis (Fig. 5): pTau first accumulates in CR-expressing ADn neurons of the Papez circuit. Next, pTau spreads to mammillothalamic vGLUT2+ terminals from ADn neuron dendrites. Myelinated axons of the ADn neurons transfer pTau to vGLUT2+ terminals in the TRN and to cortical target areas. A lack of pTau+ cell bodies in the TRN (*24*) at any stage suggests that the predominantly GABAergic cell population is ‘resistant’ to the spread of pTau from vGLUT2+ boutons, whereas cortical neurons postsynaptic to ADn neurons are likely to be vulnerable. Note we cannot currently rule out pTau spreading retrogradely from the cortex into ADn vGLUT2+ terminals, then from ADn dendrites to mammillothalamic terminals at early stages (Fig. 5). However, the very early appearance of pTau+ neurons in the ADn prior to cortical neurons makes this route unlikely. Anterograde spread of pTau from the cortex to the ADn is also unlikely due to the lack of pTau in small corticothalamic terminals at the middle stage (Braak stage III, prior to Alzheimer’s disease) or even in the late stage (Alzheimer’s disease). The vGLUT2+ mammillothalamic terminals are especially vulnerable due to a nearly 3-fold increase of pTau in these boutons from the middle stage to late stage. These boutons are unusual due to the accumulation of double-walled vesicles that may represent a type of autophagic/secretory organelle, and the presence of puncta adherentia, which contain intercellular adhesion proteins such as nectins. The puncta adherentia may even be required for the transfer of pTau (Fig. 5), and are located at other sites of potential pTau propagation including between mossy fibers of dentate granule cells and CA3 pyramidal neurons (*55, 56*). In contrast, puncta adherentia are lacking in the corticothalamic terminals in the ADn, which did not contain pTau.

Data supporting the transsynaptic spread of Tau has been obtained in animal models (*57–60*) and humans (*61, 62*), with presynaptic glutamate release inducing a rise in extracellular Tau (*63*). However, transsynaptic spread has not been previously demonstrated in the human brain at the subcellular level in well-preserved tissue. As well as ‘prion-like’ spread (*64*), Tau may also be released extracellularly (e.g. in vesicles) under certain conditions (*65, 66*). The large vGLUT2+ terminals packed with double-walled vesicles resemble those found at a lower density in the cortex of Alzheimer’s disease cases (*34, 39*). Due to their prime position within the terminal, we suggest these vesicles are candidates for the transport and/or release of Tau. Alternatively, or in addition, pTau may be directly associated with vGLUT2-containing vesicles, consistent with a previous report of Tau being associated with the cytosolic surface of vGLUT2+ synaptic vesicles (*67*).

### Selective vulnerability of different cell types

Identification of affected cell types in different neurodegenerative diseases is crucial for the understanding of biochemical factors that cause susceptibility and for therapy development, as is the case for dopaminergic neuronal loss in Parkinson’s disease (*68*). Specific cell types may be more or less vulnerable due to their connectivity, metabolic demands, or a combination. Noradrenergic locus coeruleus neurons are thought to be vulnerable due to their dense cytoarchitecture and long-range axonal projections (*69*), which could also apply to the ADn. Wolframin-expressing neurons are susceptible to pTau in human EC, and in mice pTau propagates from Wolframin+ EC cells to CA1, which is linked to memory impairments (*57*). We found that the calcium-binding protein CR was associated with pTau in the thalamus as early as Braak stage 0. This may represent a vulnerable subpopulation having specific connectivity with the cortex (e.g. with the RSg or EC) and mammillothalamic tract, forming a key part of the Papez circuit, with specific conditions for propagation (e.g. via puncta adherentia). Future immunohistochemical and RNA sequencing studies may shed light on the specific molecular identity of this subpopulation. Finally, it is recognized that glial cells also accumulate pTau (*25, 27, 70*), which can originate from pTau+ cortical pyramidal cell axons (*26*). We observed a range of pTau+ astrocyte profiles even within the same case, and ‘coiled bodies’ that are likely oligodendrocytes. Future studies will establish the contribution of different glial cell types to pTau propagation, neurodegeneration, and their association with a disrupted neurogliovascular unit (*71*), which may be one of the underlying triggers for Tau pathology in highly vulnerable brain regions such as the ADn. In summary, our results demonstrate that the connectivity, synaptic specificity, and molecular profile of subpopulations of glutamatergic rostral thalamic neurons relating to spatial orientation potentially drive progression of Tau pathology in the human brain.

## Materials and Methods

### Experimental Design

The objectives of the study were to investigate the cellular and subcellular components of the Papez circuit that are vulnerable to Tau pathology, primarily within the rostral thalamus. To achieve this, we obtained *post-mortem* human tissue from a variety of disease stages. The Braak stage was used to classify the cases into three groups (early, middle, and late stages). Details of primary antibodies, reagents, and resources are listed in Tables S5-S7.

### Human samples

Human brain samples were obtained from the Department of Pathology, Szt. Borbála Hospital, Tatabánya, Hungary via the Human Brain Research Laboratory (HBL, Institute of Experimental Medicine, Hungary), the MRC London Neurodegenerative Diseases Brain Bank (KCL, King’s College London, UK), and the Queen Square Brain Bank for Neurological Disorders (QSBB, UCL, London, UK). The reported Braak Stage (*14*) for each case was based on standardized examination of the severity and extent of pTau in the cerebral cortex. Braak stage 0 is defined by a lack of Tau pathology and no detectable cognitive impairment. At stage I, pTau is observed in the EC, followed by the hippocampus at stage II. By stage III, pTau is located other areas of the medial temporal lobe, and is associated with amyloid-ß plaques and mild cognitive impairment. Alzheimer’s disease is associated with the later stages, with pTau having spread beyond the medial temporal lobe. Psychiatric conditions for the HBL cases (Table S1) were previously assessed at the Department of Psychiatry, Szt. Borbála Hospital, Tatabánya, Hungary.

Perfusion-fixed tissue blocks from the HBL were received as previously described (*69*), and are from the same collection of cases reported by Gilvesy et al. (*69*) but with different numbering. Briefly, brains were removed 2-4 h post-mortem and vertebral arteries and internal carotid were cannulated. Perfusion was carried out using physiological saline containing 0.33% heparin (1.5 L for 30 min), followed by Zamboni fixative containing 4% paraformaldehyde and ∼0.2% w/v picric acid in 0.1 M phosphate buffer (PB), pH=7.4 (4 L for 2 h). Tissue blocks were removed and post-fixed overnight in the same solution then washed and stored in 0.1 M PB with 0.05 % sodium azide (PB-Az). Experiments were performed in compliance with the 1964 Declaration of Helsinki, approved by the Regional Committee of Science and Research Ethics of Scientific Council of Health (ETT TUKEB 31443/2011/EKU and ETT TUKEB 15032/2019/EKU). Blocks were serially sectioned into 50-µm-thick coronal sections using a Leica VTS-1000 Vibratome (Leica Microsystems, Wetzlar, Germany). Free-floating sections were incubated in 20% sucrose for cryoprotection then subjected to freeze-thaw over liquid nitrogen. Other sections were not subjected to freeze-thaw. Next, sections were incubated in 1% hydrogen peroxide to reduce endogenous peroxidase activity. Sections were washed several times in 0.1 M PB then stored in PB-Az at 4°C.

Formalin-fixed paraffin-embedded (FFPE) samples were obtained from KCL (n=6 blocks) under Tissue Bank ethics approval (18/WA/0206). A microtome (Reichert-Jung, 2035) was used to prepare 5-10 μm thick sections (up to ∼300 sections per block) and transferred onto slides (Superfrost Plus) in 37°C water. A series of 10 μm tissue sections were also obtained from QSBB (n=3 blocks) under Tissue Bank ethics approval (Human Tissue Authority Licence No 12198). In preparation for immunohistochemical tests, sections were first deparaffinized in xylene (100%) and rehydrated in a descending ethanol series (100%, 95%, 70%, 50%). Masking of epitopes caused by fixation was reversed using antigen retrieval by incubating sections in 10 mM Sodium Citrate Buffer at pH 6 at 90°C for 30 min.

### Fluorescent immunohistochemistry

Perfusion-fixed sections were blocked for 45 min in 4% Bovine Serum Albumin (BSA; Sigma) and FFPE section were blocked in 10% Normal horse serum (Vector Lab), followed by a 3-day incubation in primary antibody solution (mouse anti-AT8 diluted 1:5000, rabbit anti-Calretinin diluted 1:2000; Table S5 and S6) in 0.1 PB at 4°C. Sections were washed 3 times in 0.1 M PB then incubated in secondary antibody solution (anti-mouse Alexa Fluor 488 1:1000, anti-rabbit Cy3 1:400) in 0.1 M PB for 1 hour at room temperature (RT) or overnight at 4°C. Finally, sections were mounted in Vectashield. Every experiment included controls for the method by leaving out the primary antibodies from the full protocol.

### Brightfield immunohistochemistry

For light microscopic visualization, sections were blocked in 4% Normal Goat Serum (NGS) or 4% BSA in 0.1 M PB. This was followed by incubation with primary antibodies (1-3 days): mouse anti-AT8 1:5000, rabbit anti-Calretinin 1:1000, guinea pig anti-vGLUT2 1:500, mouse anti-vGLUT2 1:8000. We tested two other antibodies that recognize different epitopes of pTau (mouse anti-PHF1 1:1000, mouse anti-CP13 1:1000; Table S5 and S6) and observed similar distributions for pTau to that of AT8. After washing in 0.1 M PB, sections were reacted with biotinylated secondary antibodies diluted in 0.1 M PB and incubated overnight at 4°C. Subsequently, sections were incubated with 1:100 Avidin+Biotin-HRP (horseradish peroxidase) complex (Vector labs) in 0.1M PB at 4°C. The vGLUT2 immunoreaction was enhanced with tyramide signal amplification (1:50; Akoya Biosciences). Next, sections were processed using 0.5 mg/ml diaminobenzidine (DAB; Sigma-Aldrich) as chromogen, 2% nickel ammonium sulphate, and 0.4% ammonium chloride in 0.1 M PB. Subsequently, hydrogen peroxide was added to a final concentration of 0.002% w/v to initiate DAB polymerization. After 12–20 min (depending on immunolabeling intensity), reactions were stopped by washing 3 ×LJ10 min in 0.1 M PB. Free-floating sections were fixed onto glass sides using chrome alum gelatin. Next, sections were incubated in xylene for 5 min and mounted in DPX (Merck).

### Initial electron microscopic assessment of brain sections

To assess the subcellular structure of thalamic sections without any immunoreactions, sections were post-fixed in glutaraldehyde (GA) fixative (2.5% GA, ∼0.2% picric acid, 4% paraformaldehyde in 0.1 M PB) for 1-2 hours. Subsequently, sections were treated with 1% OsO_4_ in 0.1 M PB washed in 0.1 M PB and in distilled water. Next, sections were incubated in 50% and 70% ethanol, then with 1% uranyl acetate dissolved in 70% ethanol. Dehydration of sections was continued in an ascending alcohol series 70%, 90%, 95%, 100%) followed by acetonitrile. Finally, sections were embedded in epoxy resin (Durpucan AMC, Fluka, Sigma-Aldrich).

### Single- and double-labeling pre-embedding immunohistochemistry for combined light and electron microscopy

Floating sections from Cases 4, 12, 17 were washed 3x in 0.9% NaCl buffered with 50 mM Tris (pH 7.4; TBS) and then blocked in 4% NGS in TBS for 45 min. Next, sections were incubated with primary antibodies in TBS for 3 days at 4°C. The following conditions were used:

1. One primary antibody (AT8) with immunogold labeling visualized by silver intensification
2. One primary antibody (vGLUT2) with peroxidase reaction
3. Two primary antibodies, followed by a silver-intensified immunogold reaction (AT8) and a peroxidase reaction (vGLUT2)
4. Method control: no primary antibody, immunogold and biotinylated secondary antibodies followed by silver-intensification and DAB treatment

Sections were subsequently rinsed 3x in TBS and blocked for 30 min at RT in 0.1 % Cold Water Fish Skin Gelatin (CWFS) solution containing 0.8% NGS diluted in TBS to reduce the non-specific binding of secondary antibodies. Sections were incubated overnight at 4°C in CWFS solution containing biotinylated secondary antibody and/or immunogold-conjugated secondary antibodies (1:200 donkey anti-mouse ultra-small immunogold (Aurion, 100.322). Sections were washed 3x in TBS and 1x in 0.1M PB followed by incubation in 2% GA diluted in 0.1 M PB to fix the secondary antibody conjugated to immunogold particles.

After repeated washes in 0.1 M PB and TBS, sections were incubated overnight at 4°C in Avidin+Biotin-HRP complex diluted in TBS. Tissues were treated with enhancement conditioning solution (ECS; Aurion) diluted 1:10 in distilled water for 3x 5 min. To visualize immunogold particles, sections were incubated in silver enhancement solution (SE-LM, Aurion) for 20 min at 20°C in the dark and then washed in ECS. Following 2x 2 min washes in distilled water and 2x 10 min washes in 0.1 M PB, peroxidase was visualized using DAB (0.5 mg/ml) as chromogen developed with 0.01% H_2_O_2_. Subsequently, sections were washed in distilled water and treated with 0.5% OsO_4_ in 0.1 M PB for 20 min on ice in the dark. To enhance contrast, sections were incubated in 1% uranyl acetate diluted in distilled water or diluted in 70% ethanol after incubation in 50% ethanol for 25 min on ice. Next, the dehydration of sections was carried out in an ascending alcohol series (50%, 70%, 90%, 95%, 100%) followed by acetonitrile, then sections were embedded in epoxy resin. After overnight incubation at RT, they were transferred onto glass slides. To polymerize epoxy resin, sections were incubated at 55°C for 2 days. Selected regions of the thalamus were cut out and re-embedded in epoxy blocks. Series of 50-70 nm-thick sections were cut with an ultramicrotome (Leica Ultracut UTC) and collected onto pioloform-coated single slot copper grids. Some sections were counterstained with lead citrate to increase contrast. Specimens were studied on a Jeol 1010 transmission electron microscope equipped with a digital GATAN Orius camera (Department of Physiology, Anatomy and Genetics, Oxford University) and on a Hitachi 7100 electron microscope with a Veleta CCD camera (Olympus Soft Imaging Solutions, Germany), (Institute of Experimental Medicine; Budapest, Hungary). Signals were observed within the same locations and also in different structures, indicating that the experiment did not produce false-positive double-labeling results.

## Data collection

Light microscopy: Samples were obtained from 18 cases (Table S1) at different anterior-posterior levels. Three thalamic and hippocampal sections/case and two RSC sections/case were assessed.

Fluorescence immunohistochemistry: Tissues were collected from 5 cases that had at least 10 pTau+ cells in the ADn (5 cases were excluded). Three sections/case were assessed, and average values are shown.

Electron microscopy: Samples were selected from 4 cases that had sufficient quality for analysis. Data were collected from one thalamus section/case, from which 3 areas were assessed. Analysis was performed by two independent individuals (B.S. and P.S.) per case; one person was blind to the Braak stages. Only synaptic structures were quantified. Electron opaque (‘dark’) boutons were excluded from the analysis due to the likely loss of antigenicity. Dark bouton frequency was as follows: Case 4, 28.57% (n=20/70); Case 17, 29.13% (n=30/103). Tissue quality prevented an unbiased synapse density quantification, because of the large spaces amongst cellular profiles; therefore, random sampling of synapses was carried out. The frequency was influenced by the size of the boutons and dendrites. Structures were followed through 4-5 serial sections (∼4.7 sections/structure). The primary magnification was 2000-3000x, then the structures were imaged and identified at 3000-6000x magnification. Background labeling for AT8 was very low. Immunopositivity of a cellular profile for AT8 was defined by the presence of at least 1 silver-gold particle (∼12.1 particles/structure on average). Bouton area was measured across 3 serial electron micrographs and the average was calculated. Boutons without distinct membranes were not included in the analysis.

## Data analysis

To delineate thalamic nuclei in digitized AT8-immunolabeled sections, adjacent CR-immunoreacted sections were aligned with the TrakEM2 plugin in Fiji. Outlines of each thalamic nucleus were defined in the CR-immunolabeled section and imported as regions of interest (ROI) into the corresponding digitized AT8-immunolabeled section, with manual alignment required for some ROIs.

The ADn was defined by the high density of CR+ cells between the MD and AV. The AV was defined by a lack of CR+ cells, located in the dorsal part of the thalamus. The nucleus reuniens could not be accurately delineated with CR; therefore, we defined the reuniens nuclear complex (RE) as the area around the third ventricle. The AM was defined by the rostral-caudal level of sections, being located at similar level to the MB. Separation of the AM and AV at this level cannot be delineated. The PVT was defined as the dorso-ventral band of CR+ cells adjacent to the midline. The MD was defined as a large nucleus lacking CR+ neurons lateral of the PVT. The TRN was identified by its net-like structure containing CR+ cells. Imaged sections were analyzed in QuPath and Python.

For quantitative analysis, pTau coverage per thalamic nucleus was detected in QuPath using an artificial neural network (ANN_MLP) pixel classifier. Only perfusion-fixed sections were assessed, as FFPE sections lacked sufficient quality. Neuronal cell bodies were counted in QuPath by manually selecting every cell to generate counts. Every case with thalamic sections was assessed.

For pTau intensity scoring, the distribution of pTau in each area was defined by the following scores: 0, lacking detectable pTau; 0.5, containing trace inclusions; 1, sparse; 2, moderate; 3, dense. Median values for early, middle, and late-stage cases are reported. Both perfusion-fixed and FFPE sections were assessed. For cortical sections, scoring was carried out blind to the case/Braak stage.

## Imaging

Glass slides were digitized using a Pannoramic MIDI II scanner (3DHISTECH; Budapest, Hungary) with a Plan-Apochromat objective lens (20x magnification, NA 0.8, lateral resolution 0.346 x 0.325 μm/pixel) and pco.edge 4.2 4MP camera. For transmitted color brightfield images, 3 focus levels were applied. Representative images were captured in CaseViewer. For confocal microscopy, an LSM 710 was used with Plan-Apochromat 40x/1.4, 63x/1.4, and 100x/1.46 objectives (ZEN 2008 5.0 or ZEN Black 14.0 software). Laser lines (solid-state 405 nm, argon 488 nm, HeNe 543 nm and HeNe 633nm) were configured with the appropriate beamsplitters. The pinhole was set to ∼1 Airy Unit for each channel. Brightfield images were acquired with a Zeiss AX10 microscope using an AxioCam HRc camera (63x/1.4 objective). For representative confocal images, maximum intensity projections were used for z-projections.

## Data and materials availability

All data, code, and materials used in the analysis will be made available. Human brain tissue is governed by materials transfer agreements with the Brain Banks and the Human Brain Research Laboratory.

## Acknowledgments

We thank Katalin Lengyel, Kriszta Faddi, Sawa Horie, and Emily Hunter for excellent technical assistance. We thank Gabor Nyiri and Balazs Hangya for advice on tissue processing and experimental design, Andras Salma for help with slide scanning, and Cecília Szekeres-Paraczky for initial screening of the cortical tissue. We are grateful to Peter Davies for Tau antibodies and Csaba Fekete for reagents. We thank Sara Hijazi for commenting on an earlier version of the manuscript. We also acknowledge the Light Microscopy Center and Human Brain Research Laboratory (National Brain Research Program, 2017-1.2.1-NKP-2017-00002) of the Institute of Experimental Medicine, the Oxford Brain Bank (supported by the Medical Research Council, Brains for Dementia Research (Alzheimer Society and Alzheimer Research UK), Autistica UK and the NIHR Oxford Biomedical Research Centre), the MRC London Neurodegenerative Diseases Brain Bank, and the Queen Square Brain Bank for Neurological Disorders. The Queen Square Brain Bank is supported by the Reta Lila Weston Institute of Neurological Studies, UCL Queen Square Institute of Neurology.

## Author contributions

Conceptualization: BS, LA, TJV. Methodology: BS, CsD, PS, LA, TJV. Software: BS. Validation: BS, CsD, TH, PG, PS, LA, TJV. Formal analysis: BS, CsD, PS, LA, TJV. Investigation: BS, CsD, TH, PS, LA, TJV. Resources: TH, PG, PS, LA, TJV. Data curation: BS, TJV. Writing – original draft: BS, TJV. Writing – review and editing: BS, CsD, PS, LA, TJV. Visualization: BS, TJV. Supervision: PS, LA, CsD, TJV. Project administration: LA, PG, TJV. Funding acquisition: PS, LA, TJV.

