## Supplementary Materials for "Early and selective subcortical Tau pathology within the human Papez circuit"

BIORXIV/2023/543738

**This PDF file includes:**

Figs. S1 to S6
Tables S1 to S7

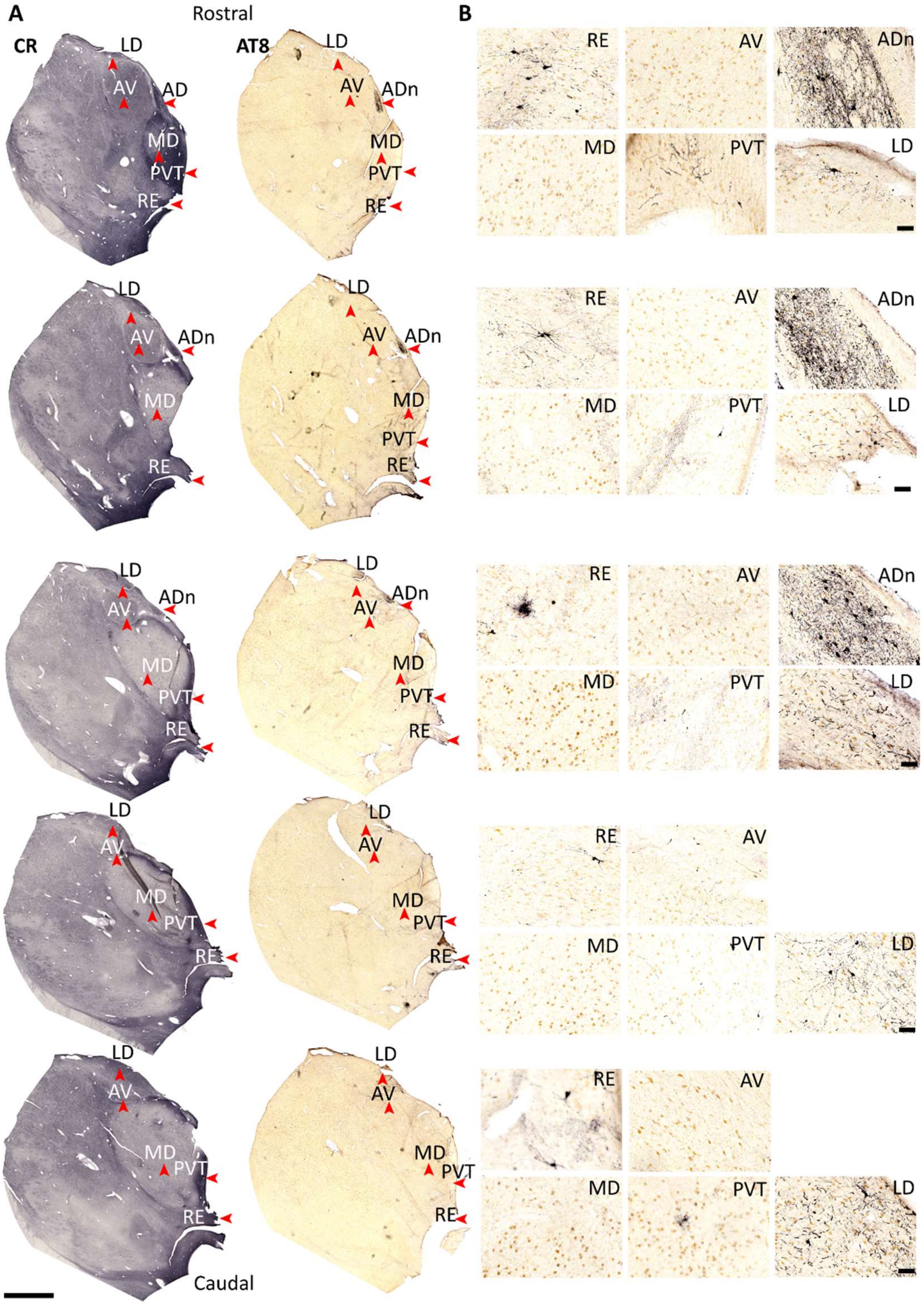


**Fig. S1. Serial sections of rostral thalamus for Braak stage III, related to Figure 1.** (**A,B**) Serial sections showing the distribution of pTau (AT8) at different rostral-caudal levels by DAB-HRP immunoreaction (nickel intensified, grey/black). Note variable pTau immunoreactivity in the laterodorsal nucleus (LD) across the rostro-caudal axis. Sections are 100 μm apart (Case 12, Braak stage III). Scale bars: 4 mm (A), 100 μm (B).

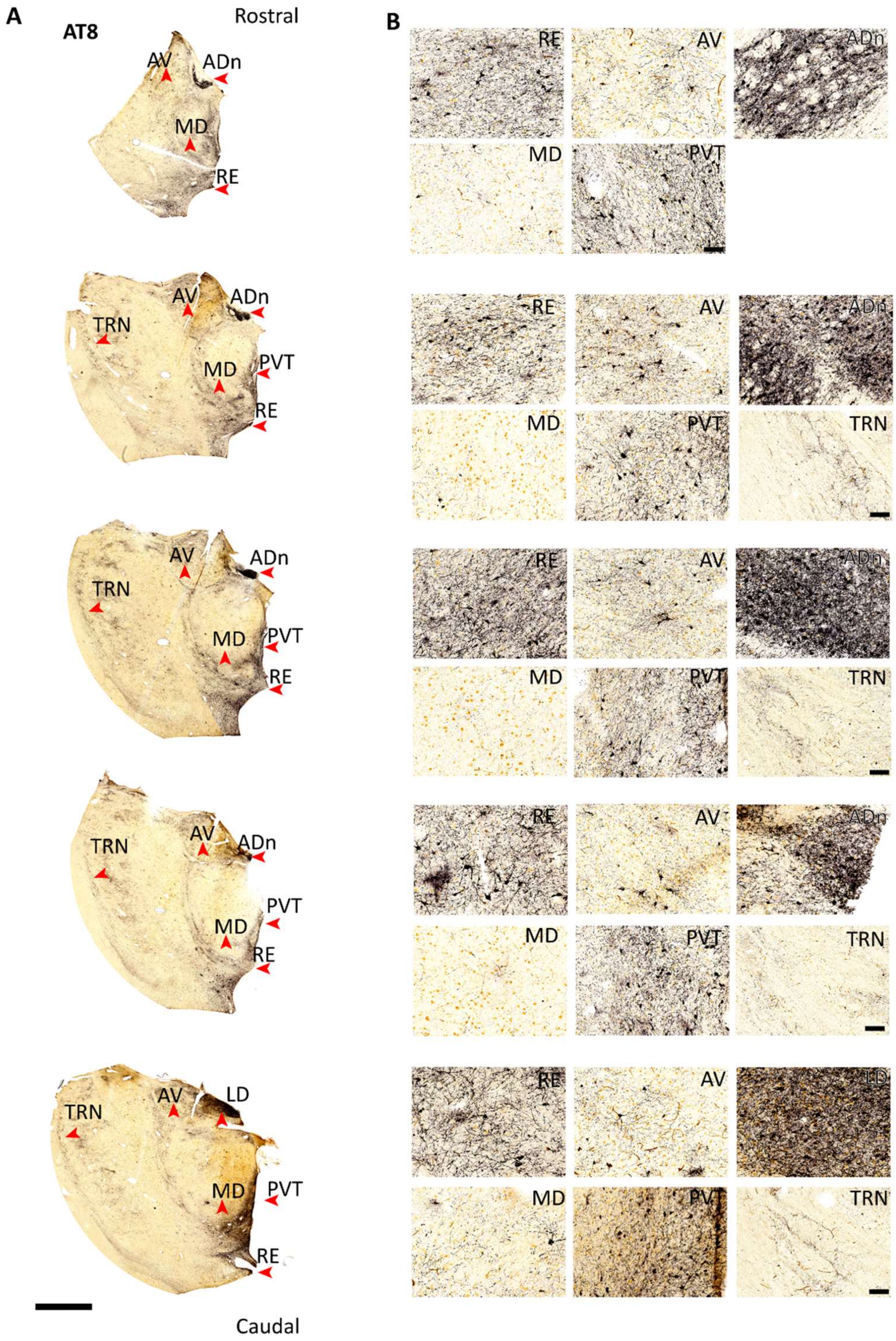


**Fig. S2. Serial sections of rostral thalamus for Braak stage VI, related to Figure 1.** (**A,B**) Serial sections showing the distribution of pTau (AT8) at different rostral-caudal levels by DAB-HRP immunoreaction (nickel intensified, grey/black). Sections are 100 µm apart (Case 17, Braak stage VI). Scale bars: 4 mm (A), 100 µm (B).

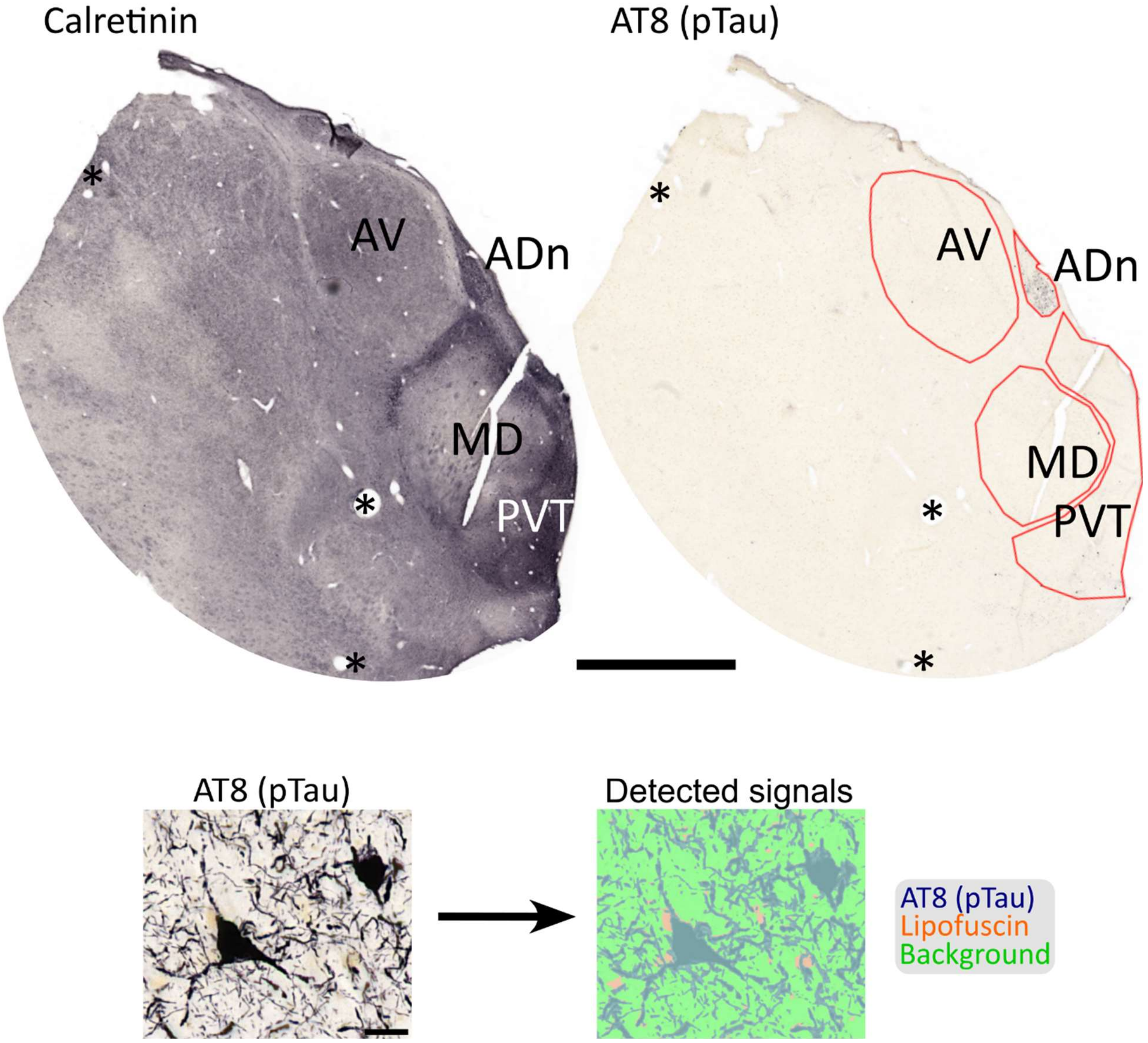


**Fig. S3. Pixel classifier, related to Figure 1.** Thalamic nuclei were delineated by CR immunoreactivity (left), followed by AT8 signal detection (right) using a pixel classifier (see Materials and Methods).

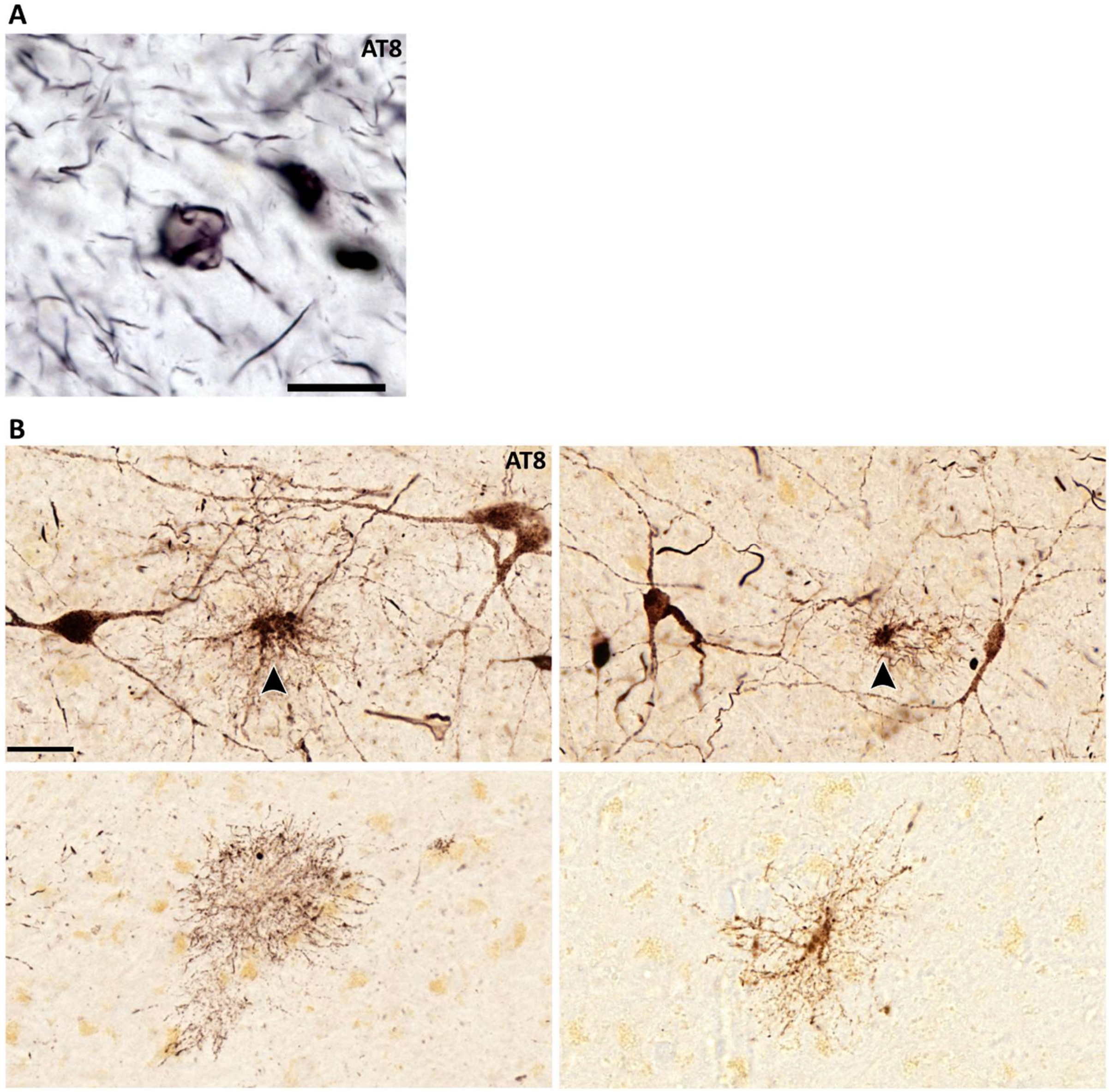


**Fig. S4. Glial cells containing pTau, related to Figure 1.** Brightfield images of glial cells, HRP-DAB immunoreaction for AT8. (**A**) A 'coiled body' in the ADn. Braak stage VI, Case 17. (**B**) A subpopulation of astrocytes were immunoreactive for pTau amongst immunopositive neurons in the posterior hypothalamic area (top panels; arrowheads) and in the AV (bottom panels). Braak stage III, Case 13. Scale bars: 10 µm (A), 40 µm (B).

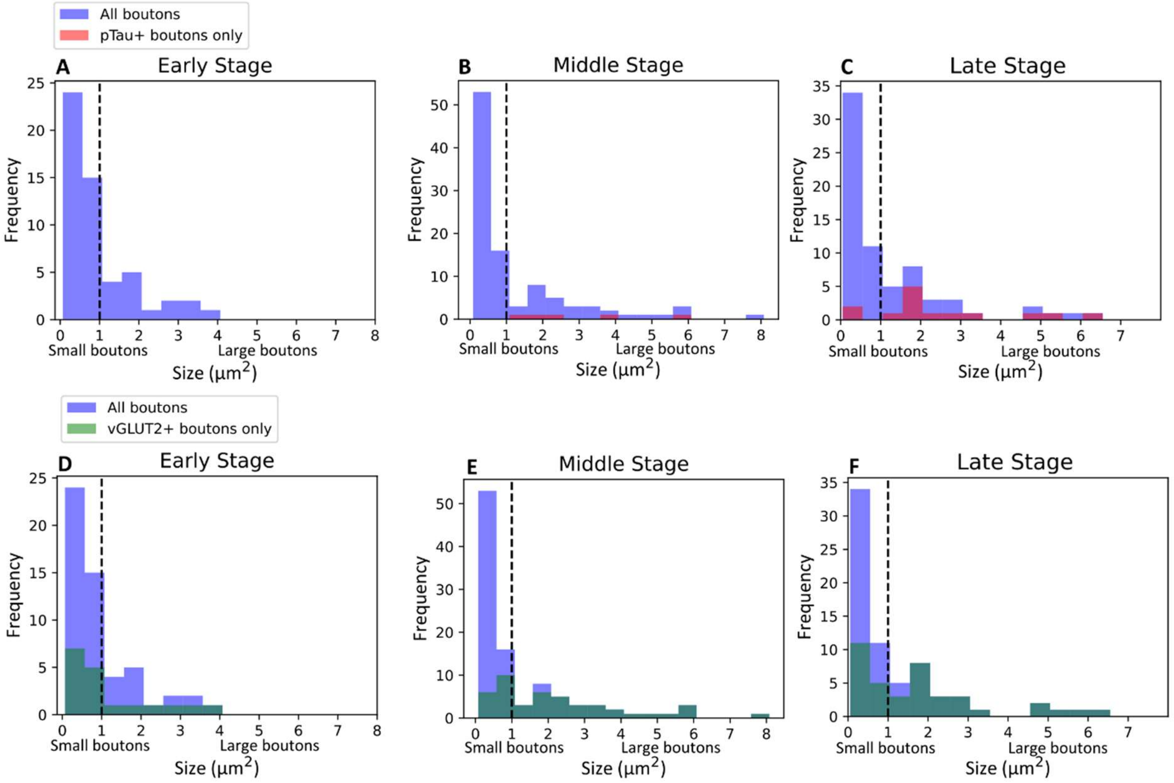


**Fig. S5. Distribution of the sizes of synaptic boutons in the ADn, related to Figures 3 and 4.** Histograms for the sizes of synaptic boutons - (blue) along with the subset that contained pTau (**A-C**) vGLUT2 (**D-F**). Early stage: Case 4; middle stage: Case 12; late stage: Case 17.

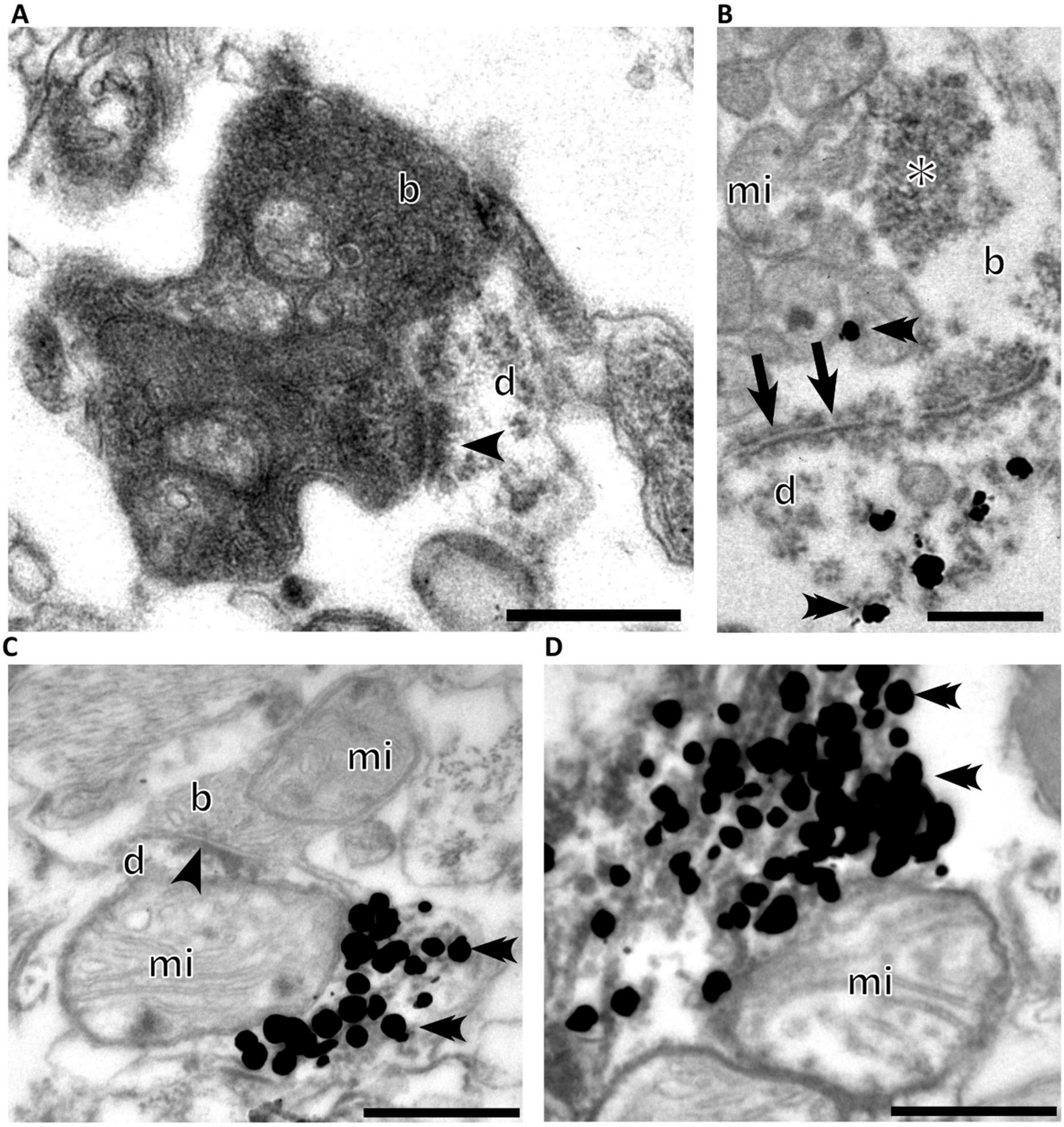


**Fig. S6. Electron micrographs of pathological structures in the ADn, related to Figures 3 and 4.** (**A**) Electron opaque ('dark') bouton (b) making a synapse (arrowhead) with a dendrite (d). Late stage, Case 17 (Braak stage VI). (**B**) Puncta adherentia (e.g. arrows) between a bouton and a dendrite. Asterisk, vGLUT2 (HRP-based DAB). Gold-silver particles (double arrowheads) recognizing pTau (AT8) were detected in both the vGLUT2+ bouton and in the dendrite. Middle stage, Case 12 (Braak stage III). (**C-D**) Dendrites containing bundles of pTau+ filaments. Mitochondria, mi. Late stage, Case 17. Scale bars: 0.5 µm.

| Case | Age (years) | Sex | Braak Stage | Thal (Aβ) phase | Clinical/ histopathological findings | PMD (h) | Sections | Source |
|---|---|---|---|---|---|---|---|---|
| Case 1 | 86 | m | 0 | na | None | 6 | FFPE | KCL |
| Case 2 | 72 | f | 0 | na | None | 36 | FFPE | QSBB |
| Case 3 | 57 | m | 0 | 0 | Paranoid schizophrenia | 2.5 | PF | HBL |
| Case 4 | 71 | f | 0 | 0 | Paranoid schizophrenia | 3 | PF | HBL |
| Case 5 | 61 | f | 0* | 4 | Paranoid schizophrenia | 4 | PF | HBL |
| Case 6 | 71 | m | I | 2 | None | 53 | FFPE | QSBB |
| Case 7 | 74 | m | I | 1 | Alzheimer's disease modified Braak stage (BrainNet Europe) 1, amyloid angiopathy | 24 | FFPE | KCL |
| Case 8 | 81 | m | I | na | Old cerebral infarct, Braak stage 1, CERAD 0 | 42 | FFPE | KCL |
| Case 9 | 72 | m | I | 1 | None | 2.5 | PF | HBL |
| Case 10 | 86 | m | III | na | Mild to moderate amyloid-beta pathology (ageing changes) | 52 | FFPE | KCL |
| Case 11 | 80 | f | III | 2 | Low probability of Alzheimer's disease | 3 | PF | HBL |
| Case 12 | 82 | f | III | 2 | Low probability of Alzheimer's disease | 2 | PF | HBL |
| Case 13 | 84 | m | III | 1 | Catatonic schizophrenia | 3 | PF | HBL |
| Case 14 | 89 | f | IV | 0 | Intermediate probability of Alzheimer's disease | 2.5 | PF | HBL |
| Case 15 | 78 | m | V | 5 | Circumscribed brain atrophy (frontotemporal dementia). Neuropathological diagnosis: Alzheimer's disease | 5 | FFPE | QSBB |
| Case 16 | 71 | m | V/VI | na | Alzheimer's disease | 5 | FFPE | KCL |
| Case 17 | 81 | f | VI | 4-5 | High probability of Alzheimer's disease | 2 | PF | HBL |
| Case 18 | 72 | m | VI | na | Alzheimer's disease | 5 | FFPE | KCL |
| Case 19** | 76 | f | VI | na | Alzheimer's disease | 2.2 | PF | HBL |

**Table S1. List of cases, related to Figures 1-4.** Braak stages, determined by independent neuropathologists not involved in this investigation. Braak stage 0 is defined as no reported (or negligible) cortical Tau pathology. *Classified as Braak Stage 0 based on available material (prior assessment by a neuropathologist unavailable). **Only cortical sections were available. Abbreviations: na, not available; PMD, *post-mortem* delay time; PF, perfusion-fixed (50-µm-thick free-floating sections); FFPE, formalin-fixed paraffin-embedded (10-µm-thick sections); CERAD, Consortium to Establish a Registry for Alzheimer's Disease; KCL, London Neurodegenerative Diseases Brain Bank, King's College London; QSBB, Queens Square Brain Bank, University College London; HBL (Human Brain Research Laboratory), Institute of Experimental Medicine, Budapest.

| Case | Stage | Cause of death | Comorbidities | Polymorphisms |
|---|---|---|---|---|
| Case 1 | 0 | Myocardial infarction | None reported | APOE 3,3 |
| Case 2 | 0 | Metastatic lung cancer | Mild arteriolar CAA, mild cerebrovascular atherosclerosis, capillary CAA | |
| Case 3 | 0 | Cardiac arrest | Mild SVD, atherosclerotic heart disease, bronchitis, emphysema | |
| Case 4 | 0 | Respiratory arrest | Myocardial degeneration, heart failure, asthma, bronchitis, emphysema | |
| Case 5 | 0 | Heart failure | Chronic ischemic heart disease, hypertension, atherosclerosis, gastroenteritis, colitis, paralytic ileus, obesity | |
| Case 6 | I | Bronchopneumonia | Mild non-amyloid SVD, mild cerebrovascular atherosclerosis | na |
| Case 7 | I | Respiratory arrest; cancer | Mild arteriolar CAA | APOE 3/3 |
| Case 8 | I | Duodenal cancer | Stroke | APOE 3,3 |
| Case 9 | I | Respiratory arrest | Mild SVD, Atherosclerosis unsp, Subsequent myocardial infarct anterior wall, Abscess of lung with pneumonia, Hydrothorax, Acute bronchitis unsp., Hepatomegaly | |
| Case 10 | III | Heart failure | None reported | APOE 3/3 |
| Case 11 | III | Functional intestinal disorder | Atherosclerotic heart disease of native coronary artery without angina pectoris, Hypertonia, Dilated cardiomyopathy | |
| Case 12 | III | Myocardial infarction | Atherosclerosis, hypertonia, heart disease | |
| Case 13 | III | Respiratory arrest | Catatonic schizophrenia, Sick sinus syndrome, Presence of cardiac pacemaker, Emphysema unsp., Disease of pancreas, Comorbidity: vascularis encephalopathy | |
| Case 14 | IV | na | ARTAG, Tau-positive grains | |
| Case 15 | V | Rectal cancer | Mild non-amyloid SVD, capillary CAA, moderate arteriolar Aß-CAA, limbic Lewy body disease, TDP43opathy | na |
| Case 16 | V/VI | Gastrointestinal bleeding | Not reported | APOE 3,4 |
| Case 17 | VI | na | Moderate SVD, possible TDP43. | |
| Case 18 | VI | na | CAA | APOE 3/3 |
| Case 19 | VI | Cardiac failure | Arteriosclerosis, ischemic heart disease, hypertonia | na |

**Table S2. Additional information on cases, related to Figures 1-4.** Abbreviations: SVD, small vessel disease. ARTAG, aging-related Tau astrogliopathy. CAA, cerebral amyloid angiopathy. Not available, na.

| Case | Braak stage | Nucleus | pTau coverage (%) | pTau cell frequency (cell/mm$^2$) |
|---|---|---|---|---|
| 4 | 0 | AV | 0.07 | 0 |
| | | ADn | 0.84 | 2.58 |
| | | MD | 0.06 | 0 |
| | | PVT | 0.66 | 0.32 |
| | | TRN | 0.08 | 0 |
| 5 | 0 | AV | 0.09 | 0 |
| | | ADn | 0.28 | 0 |
| | | MD | 0.05 | 0 |
| | | PVT | 0.11 | 0.04 |
| | | TRN | 0.07 | 0 |
| 11 | III | AV | 0.08 | 0.11 |
| | | ADn | 3.31 | 4.97 |
| | | MD | 0.07 | 0.02 |
| | | PVT | 0.13 | 0.05 |
| | | TRN | 0.04 | 0 |
| 12 | III | AV | 0.17 | 0.03 |
| | | ADn | 12.14 | 15.13 |
| | | MD | 0.19 | 0 |
| | | PVT | 0.40 | 0.47 |
| | | TRN | 0.09 | 0 |
| 13 | III | AV | 0.31 | 0.32 |
| | | ADn | 4.52 | 14.84 |
| | | MD | 0.30 | 0.93 |
| | | PVT | 1.99 | 5.2 |
| | | TRN | 0.19 | 0.03 |
| 14 | III/IV | AV | 0.18 | 0.14 |
| | | ADn | 14.20 | 19.59 |
| | | MD | - | - |
| | | PVT | 1.34 | 4.58 |
| | | TRN | 0.07 | 0 |
| 17 | VI | AV | 6.28 | 14.49 |
| | | ADn | 36.31 | 0 |
| | | MD | 1.62 | 2.84 |
| | | PVT | 11.62 | 29.13 |
| | | TRN | 2.98 | 0 |

**Table S3. Quantification of pTau coverage and immunopositive cell counts, related to Figures 1, 2 and S3.** Pixel-classifier quantification of pTau coverage and cell counts from perfusion-fixed tissue.

| Location Stage: | 0 | I | III | IV | V/VI | VI |
|---|---|---|---|---|---|---|
| ADn | 0(1), 1(2),2(1) | 1 (2) | 1(1), 2(3) | 2(1) | 3(2) | 3 (1) |
| AV | 0(5) | 0 (3) | 0 (3), 1(1) | 1(1) | 2(1) | 2 (1) |
| LD | 0(1) | 0 (1) | 0(1), 1(1) | na | 3(1) | 2(1), 3(1) |
| MD | 0(4) | 0(3) | 0 (3) 1(1) | 0(1) | 1(1) | 1(2) |
| PVT | 0(3), 0.5(2) | 0(2) | 0(1), 0.5(1), 1(1), 2(1) | 2(1) | 2(1), 3(1) | 2(1), 3(1) |
| TRN* | 0(2), 0.5(3) | 0(2) | 0(1), 1(4) | 0(1) | 2(2) | 2 (2) |
| RE | 0(3), 0.5(1) | 0(2) | 0(1), 1(3) | 1 (1) | 2(1) | 2 (2) |
| DG | 0(3) | na | 0(1),0.5(1), 1(1) | na | na | 2 (1), 3(1) |
| CA3 | 0(1), 0.5(1), 1(1) | na | 1(1), 2(1), 3 (1) | na | na | 3 (2) |
| CA2 | 0 (1),0.5 (1) 1(1) | na | 1(1), 2 (1), 3 (1) | na | na | 3 (2) |
| CA1 | 0(2), 1(1) | na | 2(1), 3(2) | na | na | 3 (2) |
| Prosubiculum | 0(2), 2 (1) | na | 2(2), 3(3) | na | na | 3 (2) |
| Subiculum | 0(2), 1(1) | na | 1(2), 2(1) | na | na | 3 (2) |
| Presubiculum | 0(2),0.5(1) | na | 0(1), 1(2) | na | na | 3 (2) |
| Parasubiculum | 0(2), 1(1) | na | 1(2), 3(1) | na | na | 3 (2) |
| Entorhinal area | 0(1), 1(1) 2(1) | na | 3(3) | na | na | 3 (1) |
| Retrosplenial area | 0(1), 1(1) | na | 3(3) | na | na | 2(1), 3 (1) |
| Cingulate area | 0(2) | na | 1(3) | na | na | 3 (1) |

**Table S4. Intensity scoring for pTau in the rostral thalamus and cerebral cortex, related to Figure 1.** The distribution of pTau in each area was defined by the following scores: 0, lacking detectable pTau; 0.5, containing trace inclusions; 1, sparse; 2, moderate; 3, dense. Numbers in parentheses show the number of cases with that score out of the total in the given stage. The TRN contained a few pTau immunopositive axons even when score 0 was given (asterisk).

| Primary antibody | Source | Identifier |
|---|---|---|
| Mouse anti-AT8 | Thermo Fisher Scientific | Cat# MN1020; RRID: AB_223647 |
| Rabbit anti-Calretinin | Swant | Cat# 7699/3H; RRID: AB_10000321 |
| Mouse anti-Vesicular glutamate transporter 2 | Sigma Aldrich | Cat# MAB5504; RRID: AB_2187552 |
| Guinea pig anti-Vesicular glutamate transporter 2 | Synaptic Systems | Cat# 135 404; RRID: AB_887884 |
| Mouse anti-CP13 | Dr Peter Davies | |
| Mouse anti-PHF-1 | Dr Peter Davies | |

**Table S5. Primary antibodies (I/II).**

| Primary antibody (repeated) | Concentration and dilution | Epitope | Specificity information |
|---|---|---|---|
| Mouse anti-AT8 | 200 µg/mL, 1:2500 for DAB 1:5000 for IF | Phosphorylation at S202 and T205 residues of human Tau | Immunoblot (Goedert et al., 1995) |
| Rabbit anti-Calretinin | Antiserum, 1:1000 for DAB 1:2000 for IF | Polyclonal, recombinant human calretinin with 6-his tag | Western blot supplied by Swant. No signal in knockout animals |
| Mouse anti-vGLUT2 | Purified, 1:500 for DAB | Recombinant rat vGLUT2 amino acids 510-582 | Similar to immunoreactivity characterized in mouse hippocampus by Herzog et al. 2006 J. Neurochem. |
| Guinea pig anti-vGLUT2 | Antiserum, 1:8000 for DAB with Tyramide | Recombinant protein from rat vGLUT2 | Evaluated by Western Blot on Mouse brain lysates (Sigma Aldrich) |
| Mouse anti-CP13 | Cell culture supernatant, 1:1000 for DAB | Detects Tau phosphorylated at S202. | Tau knock out mice |
| Mouse anti-PHF-1 | Cell culture supernatant, 1:1000 for DAB | Around S396 and S404 phosphorylated sites | Tau knock out mice |

**Table S6. Primary antibodies (II/II).**

| **Chemicals** | | |
|---|---|---|
| Normal goat serum | Vector Laboratories | Cat# S-1000 |
| Normal horse serum | Vector Laboratories | Cat# S-2000 |
| Cold Water Fish Skin Gelatin | Aurion | Cat# 900.033 |
| Enhancement conditioning solution (ECS) | Aurion | Cat# 500.055 |
| Silver enhancement solution SE-LM | Aurion | Cat# 500.022 |
| Diaminobenzidine (DAB) | Sigma-Aldrich | Cat# D5637-1G |
| Chromium(III) potassium sulfate dodecahydrate | Sigma-Aldrich | Cat# 243361 |
| Gelatin from bovine skin | Sigma-Aldrich | Cat# G9391 |
| Acetonitrile | VWR Chemicals | Cat# 83657.32 |
| DPX Mountant | Merck | Cat# 6522 |
| Donkey anti-mouse Alexa Fluor 488 | Thermo Fisher Scientific | Cat# A-21202 |
| Donkey anti-rabbit Cy3 | Jackson Immuno Research | Cat# 711-165-152 |
| Biotinylated Goat anti-mouse | Vector Laboratories | Cat# BA-9200-1.5 |
| Goat-anti-Mouse IgG (H&L) Ultra Small | Aurion | Cat# 800.022 |
| **Critical commercial assays** | | |
| Vectastain ABC Elite kit | Vector Laboratories | Cat# PK6100; RRID: AB_2336819 |
| Vectashield Antifade Mounting Medium | Vector Laboratories | Cat# H-1000; RRID: AB_2336789 |
| TSA Biotin Reagent Pack | Akoya Biosciences | Cat# SAT700001EA |
| **Software and algorithms** | | |
| Fiji | ImageJ | https://imagej.net/software/fiji/ |
| Zen 2008, Zen Black, Zen Blue, | Zeiss | www.zeiss.co.uk |
| Python | | https://www.python.org/ |
| CaseViewer | 3DHISTECH | https://www.3dhistech.com/solutions/caseviewer/ |
| QuPath | Bankhead et al. | https://qupath.github.io/ |
| TrakEM2 | Cardona et al. | https://www.ini.uzh.ch/~acardona/trakem2.html |

**Table S7. Other reagents and resources.**
